# Minimal synthetic enhancers reveal control of the probability of transcriptional engagement and its timing by a morphogen gradient

**DOI:** 10.1101/2021.07.10.451524

**Authors:** Armando Reimer, Simon Alamos, Clay Westrum, Meghan A. Turner, Paul Talledo, Jiaxi Zhao, Hernan G Garcia

## Abstract

How enhancers interpret morphogen gradients to generate spatial patterns of gene expression is a central question in developmental biology. Although recent studies have begun to elucidate that enhancers can dictate whether, when, and at what rate a promoter will engage in transcription, the complexity of endogenous enhancers calls for theoretical models with too many free parameters to quantitatively dissect these regulatory strategies. To overcome this limitation, we established a minimal synthetic enhancer system in embryos of the fruit 2y *Drosophila melanogaster*. Here, a gradient of the Dorsal activator is read by a single Dorsal binding site. By quantifying transcriptional activity using live imaging, our experiments revealed that this single Dorsal binding site is capable of regulating whether promoters engage in transcription in a Dorsal concentration-specific manner. By modulating binding-site affinity, we determined that a gene’s decision to engage in transcription and its transcriptional onset time can be explained by a simple theoretical model where the promoter has to traverse multiple kinetic barriers before transcription can ensue. The experimental platform developed here pushes the boundaries of live-imaging in studying gene regulation in the early embryo by enabling the quantification of the transcriptional activity driven by a single transcription factor binding site, and making it possible to build more complex enhancers from the ground up in the context of a dialogue between theory and experiment.

## 1 Introduction

The adoption of distinct cellular identities in multicellular organisms relies on the formation of spatial gene expression domains driven, in large part, by transcriptional regulatory programs. The positional information giving rise to these mRNA patterns is typically provided by transcription factor gradients (Fig. 1A) whose concentrations are interpreted by enhancer DNA sequences that, in turn, regulate transcription of developmental genes (***Wolpert, 1969***; ***Briscoe and Small, 2015***). A long-standing goal in quantitative developmental biology is to precisely predict gene expression from knowledge of the DNA regulatory sequence and morphogen concentration (***Garcia et al., 2020***; ***Vincent et al., 2016***). Achieving this predictive understanding requires theoretical models that calculate how DNA sequence dictates the functional relation between input morphogen concentration and output transcriptional activity, and calls for testing these predictions by measuring input-output functions (***Garcia et al., 2020***). Precise genetic manipulations (***Venken and Bellen, 2005***; ***Bier et al., 2018***) and powerful imaging technologies (***Gregor et al., 2005***; ***Garcia et al., 2013***; ***Mir et al., 2017***) have rendered the early embryo of the fruit 2y *Drosophila melanogaster* (*Drosophila*) a prime model system for quantitatively dissecting these input-output functions in development.

**Figure 1.**
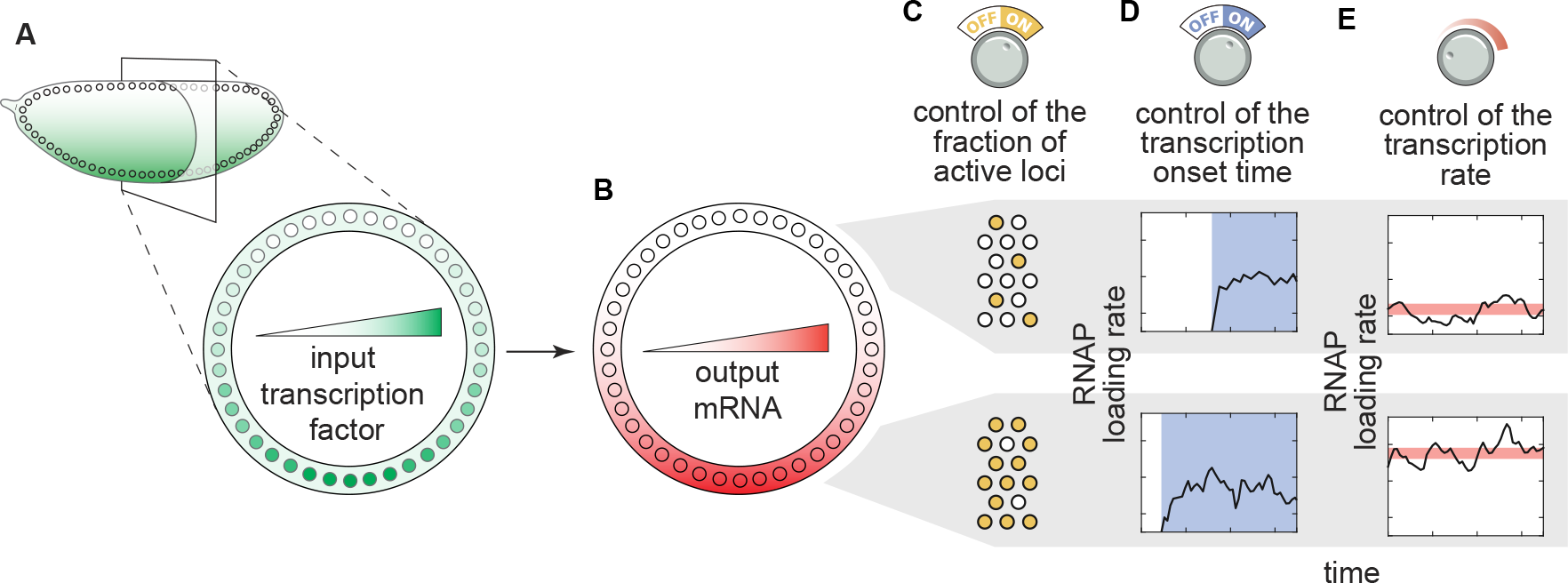
Transcriptional regulatory strategies of enhancers in response to transcription factor concentration gradients. **(A)**A *Drosophila* embryo with a transcription factor gradient along its dorsoventral axis. **(B)** This input transcription factor dictates the emergence of output gene-expression patterns by controlling a combination of three enhancer regulatory ‘knobs’: **(C)** the probability of loci becoming transcriptionally active, **(D)** the transcriptional onset time, and **(E)** the mean transcription rate of active loci. (RNAP, RNA polymerase II).

In recent years, several studies have reported that *Drosophila* enhancers can control various, potentially independent aspects of transcriptional dynamics in early embryonic development (Fig. 1; ***Lucas et al.*** (***2013***); ***Garcia et al.*** (***2013***); ***Fukaya et al.*** (***2016a***); ***Lammers et al.*** (***2020***); ***Fuqua et al.*** (***2020***); ***Eck et al.*** (***2020***); ***Berrocal et al.*** (***2020***); ***Fukaya*** (***2021***); ***Harden et al.*** (***2021***)). First, for a given gene, a fraction of loci remain transcriptionally inactive throughout entire mitotic cycles in development, even when exposed to the same activator concentration as active loci (Fig. 1B)—a behavior usually quantified through the fraction of active nuclei or loci. This stochastic decision for a locus to become active is a ubiquitous and potentially important regulatory feature for shaping gene-expression patterns in the embryo (***Garcia et al., 2013***; ***Dufourt et al., 2018***; ***Lammers et al., 2020***; ***Harden et al., 2021***). However, it remains unclear whether this feature constitutes a regulatory ‘knob’ or whether inactive loci are artifacts of experimental detection thresholds. Second, the timing of transcription onset (and cessation, which is not addressed in the present investigation) can also be controlled by input transcription-factor dynamics (Fig. 1C; ***Desponds et al.*** (***2016***); ***Tran et al.*** (***2018***); ***Dufourt et al.*** (***2018***); ***Eck et al.*** (***2020***); ***Lammers et al.*** (***2020***); ***Desponds et al.*** (***2020***); ***Harden et al.*** (***2021***)). Finally, the rate of transcriptional initiation in active loci is under regulatory control (Fig. 1D) and has been the focus of most studies to date (e.g., ***Garcia et al.*** (***2013***); ***Fukaya et al.*** (***2016b***); ***Park et al.*** (***2019***); ***Lammers et al.*** (***2020***); ***Berrocal et al.*** (***2020***); ***Fukaya*** (***2021***)). Thus, multiple regulatory strategies together realize gene-expression patterns in space and time (Fig. 1E).

Intense theoretical scrutiny (*Desponds et al., 2016*; *Fakhouri et al., 2010*; *Sayal et al., 2016*; *Estrada et al., 2016*; *Scholes et al., 2017*; *Dufourt et al., 2018*; *Park et al., 2019*; *Eck et al., 2020*; *Cheng et al., 2021*) has generated a compelling hypothesis: that the regulation of transcriptional dynamics can be separated into two stages. First, a promoter must pass through a series of kinetic barriers consisting of reactions catalyzed by transcription factors in order for for loci to engage in transcription. Previous analyses of the mean and distribution in transcriptional onset times have suggested that the number of inactive promoter states can range from one to three (*Dufourt et al., 2018*; *Eck et al., 2020*; *Harden et al., 2021*). These reactions could be associated with, for example, the stepwise unwrapping of DNA from nucleosomes (*Desponds et al., 2016*; *Dufourt et al., 2018*; *Eck et al., 2020*) and/or the sequential recruitment of general transcriptional cofactors (*Zhou et al., 1998*). Second, after initial promoter activation, the rate of mRNA production is proportional to the probability of finding RNA polymerase II (RNAP) bound to the promoter. Statistical mechanical (also called thermodynamic) models have been used to calculate this probability of finding RNAP bound to the promoter, and have successfully use to predict mRNA production rates in bacteria (*Razo-Mejia et al., 2018*). However, whether they can be applied to the more complex context of eukaryotic transcriptional regulation—let alone to the dynamical processes of cellular decision-making in development—is still an open question (*Polach and Widom, 1995*; *Schulze and Wallrath, 2006*; *Lam et al., 2008*; *Li et al., 2008*; *Kim and O’Shea, 2008*; *Levine, 2010*; *Fussner et al., 2011*; *Bai et al., 2011*; *Li et al., 2014*; *Hansen and O’Shea, 2015*; *Estrada et al., 2016*; *Li and Eisen, 2018*; *Park et al., 2019*; *Eck et al., 2020*).

One of the main challenges to systematically testing these models is the complexity of endogenous regulatory regions (***Fakhouri et al., 2010***; ***Foo et al., 2014***; ***Sayal et al., 2016***; ***Dufourt et al., 2018***; ***Park et al., 2019***; ***Eck et al., 2020***). Because endogenous enhancers contain multiple binding sites for different transcription factors, accounting for these sites and their interactions leads to a combinatorial explosion of model parameters (***Garcia et al., 2016***, ***2020***); determining the values of these parameters from simple experiments constitutes a computational—and conceptual—challenge (***Vincent et al., 2016***; ***Garcia et al., 2016***, ***2020***). To render complex transcriptional regulatory systems tractable to theory, minimal synthetic enhancers have been engineered in bacteria (***Garcia and Phillips, 2011***; ***Brewster et al., 2014***; ***Razo-Mejia et al., 2018***; ***Phillips et al., 2019***), eukaryotic cells (***Popp et al., 2020***), and developing organisms (***Fakhouri et al., 2010***; ***Sayal et al., 2016***). In such experiments, a short, synthetic DNA sequence with only one to a few binding sites for a single transcription factor drives the expression of a reporter gene. Measuring the concentration of the transcription-factor input and reporter mRNA output makes it possible to test models of transcriptional regulation and to infer molecular parameters that can be used to predict the behavior of more complex regulatory architectures (***Phillips et al., 2019***).

Here we sought to use synthetic minimal enhancers to challenge our integrated model of transcriptional control using the dorsoventral patterning system in *Drosophila* embryos, in which a concentration gradient of the Dorsal transcription factor specifies spatial domains of transcrip- tion, as a case study. To test the integrated model of transcriptional dynamics (Fig. 2A,B), we performed simultaneous quantitative live-cell measurements of Dorsal concentration (input) and transcription (output) driven by minimal synthetic Dorsal-dependent enhancers in single nuclei. By repurposing the *parS*-ParB DNA labeling technology (***Germier et al., 2017***; ***Chen et al., 2018***) to quantify transcriptional activity independent of RNA detection, we determined that the inactive loci described by our model constitute a distinct transcriptional state under regulatory control and are not the result of detection artifacts. Further, our theoretical model predicted how, through the Dorsal-mediated catalysis of reactions prior to transcriptional onset, regulatory architecture dictates both the transcriptional onset time and the fraction of active loci. Finally, once promoters turn on, we found that our measurements are compatible with an equilibrium model. Thus, the present investigation provides quantitative evidence supporting a unified model of transcriptional regulation in eukaryotes that accounts for whether loci become transcriptionally active, when this activity ensues, and, once transcription ensues, at what rate nascent RNA molecules are produced. More generally, our work demonstrates the feasibility of using minimal synthetic enhancers to engage in a dialogue between theory and experiment in the context of transcriptional control in development.

**Figure 2.**
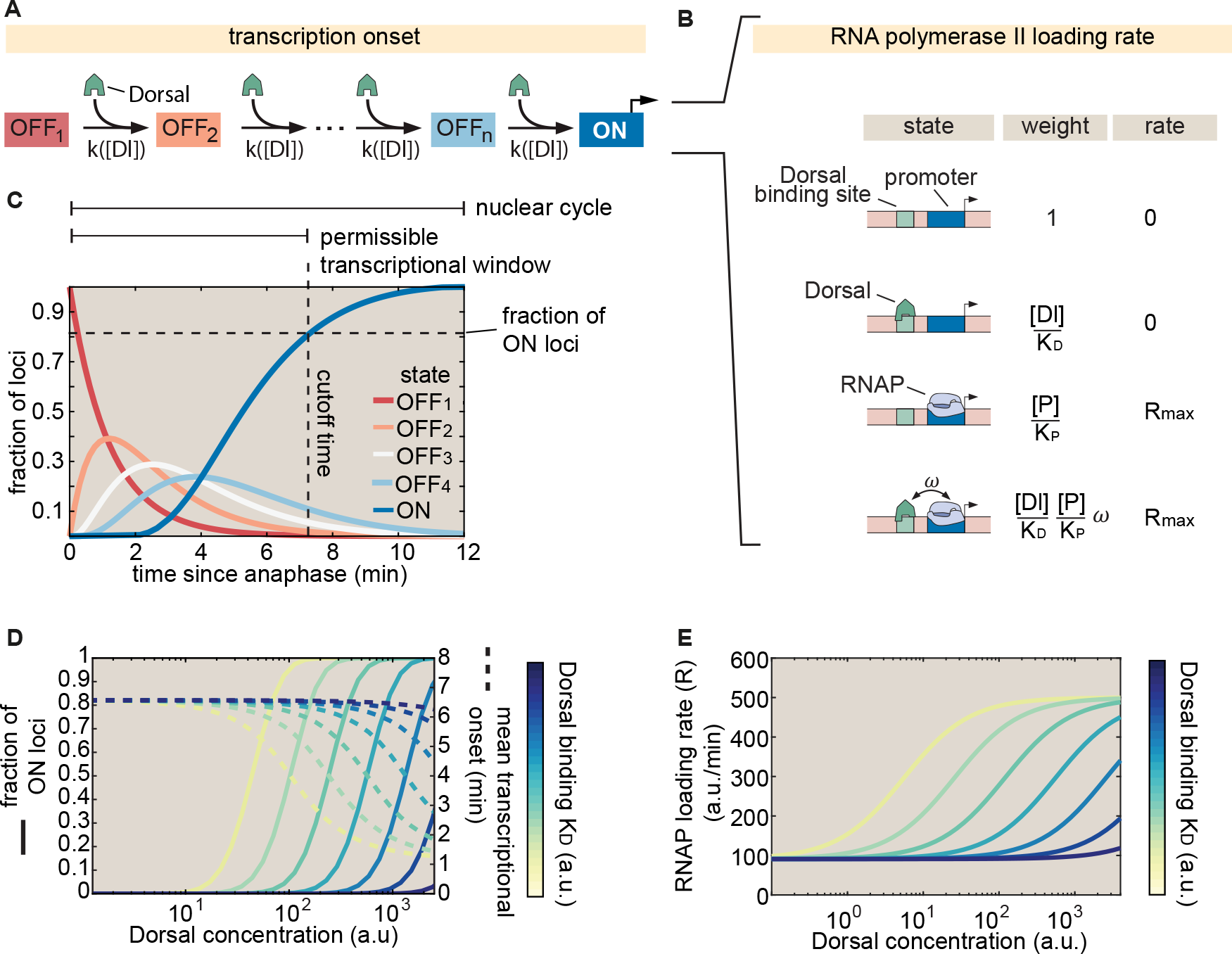
Integrated kinetic and thermodynamic model of simple activation by Dorsal. **(A)** The promoter undergoes kinetic transitions from transcriptionally inactive states (OFF_1_ to OFF*_n_*) to an active state (ON) with Dorsal accelerating the transition rate, *k*, by a factor proportional to the Dorsal occupancy at the promoter. **(B)** Thermodynamic states and weights for the simple activator model. The probability of 1nding RNAP bound to the promoter can be calculated from the statistical weights associated with all possible occupancy states of the enhancer-promoter system. **(C)** Visualization of a particular solution of the kinetic scheme from (A) showing the probability of 1nding a given locus in each of the states for an illustrative, representative set of parameters ([*Dl*] = 1000 a.u., *K_D_* = 1000 a.u., *c* = 10/min, *n* = 4 states, and 7 min nuclear cycle duration). The predicted fraction of active loci (dashed horizontal line) is calculated as the probability of being in the ON state by the end of the permissible time window (dashed vertical line) that is determined by mitotic repression. **(D)** Predictions for the fraction of active loci (solid lines plotted against the left y-axis) and mean transcriptional onset times (dashed lines plotted against the right y-axis) as a function of Dorsal concentration for different, illustrative values of the Dorsal dissociation constant *K_D_* . **(E)** Rate of mRNA production across active loci as a function of Dorsal concentration for different values of *K_D_* based on the model in (B) (*R_max_* = 1000 a.u., Dorsal *K_D_* ranging from 10 a.u. to 10^5^ a.u., *w* = 10, [*P* ]/[*K_P_* ] = 0.1).

## 2 Results

### 2.1 An integrated model of transcriptional dynamics driven by a single activator binding site

To probe the transcriptional regulatory strategies (Fig. 1) of a minimal synthetic enhancer, we posit a theoretical model that predicts the fraction of loci that will become active, their transcriptional onset time, and RNAP loading dynamics once transcription ensues. Specifically, we consider a simplified case in which only one activator is present and can only bind to one site only a few base pairs away from the promoter (Fig. 2).

In order to explain the transcriptional onset dynamics of a locus and the probability of loci becoming active, we invoke recent experiments leading to a ‘kinetic barrier’ model (***Desponds et al., 2016***; ***Dufourt et al., 2018***; ***Eck et al., 2020***) proposing that, after exiting mitosis, all promoters are in an inactive state. In this state, labeled as ‘OFF_1_ ’ in Figure 2A, transcription is not possible. Promoters must then traverse a series of distinct inactive states (labeled ‘OFF_2_ ’ to ‘OFF*_n_*’ in Fig. 2A) before reaching an active state in which transcription proceeds (labeled ON in Fig. 2A).

The temporal evolution of the enhancer-promoter system as it traverses the states shown in Figure 2A can be simulated by computing the probability that the promoter occupies each state. Here, the transition rates between states, *k*, determines how the states probability spreads from the initial condition where the promoter is in state OFF_1_ to the active state as time passes (see Section S1.1 for details).

We propose that a transcriptional activator such as Dorsal can catalyze the transition between states in an affinity-dependent manner via binding to its cognate site in the enhancer. Because we assume that Dorsal binding and unbinding is faster than the transition rate *k*, we posit that *k* is a linear function of the equilibrium Dorsal occupancy at the enhancer such that

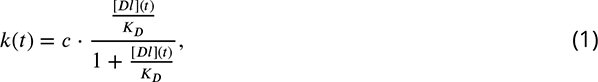

where *c* is a rate constant, [*Dl*](*t*) is the Dorsal concentration at time *t*, and *K_D_* is the Dorsal-DNA dissociation constant.

Because Dorsal is time-varying, the model cannot be solved analytically. As a result, we numerically calculated the probability of the promoter being in each state as a function of time using a particular set of model parameters (see details in Section S1 .1). As seen in Figure 2C, because individual loci must traverse a sequence of intermediate states before reaching the ON state, this model introduces a delay in activation.

This kinetic barrier model accounts for loci that never transcribe during the nuclear cycle. Specifically, if nuclear cycles lasted indefinitely, all promoters would eventually reach the ON state as shown in Figure 2C. However, due to the rapid mitotic cycles that characterize early embryonic development in *Drosophila*, this duration is limited: transcription cannot initiate during mitosis and thus is only permissible during a time window within interphase (Fig. 2C, vertical dashed line; ***Shermoen and O’Farrell*** (***1991***); ***Garcia et al.*** (***2013***); ***Eck et al.*** (***2020***)). Consequently, if the time it takes a promoter to reach the ON state is longer than the duration of this window, then this hypothetical promoter will not initiate transcription at all during the nuclear cycle (Fig. 2C, horizontal dashed line).

The kinetic barrier model can be used to predict two of the three regulatory strategies, fraction of active loci and transcription onset times, that we aim to dissect quantitatively (Fig. 1). First, the model predicts how the fraction of active loci is determined by Dorsal nuclear concentration and binding affinity (Fig. 2D, left y-axis). Second, this same model calculates the mean transcriptional onset time of those loci that turn on as a function of these same Dorsal parameters (Fig. 2D, right y-axis).

To model a locus once it is active, we follow ***Eck et al.*** (***2020***) and propose a simple thermodynamic model (***Bintu et al., 2005b***,a) that assumes that the RNAP loading rate, *R*, is proportional to the probability of finding RNAP bound to the promoter *p_bound_* , such that

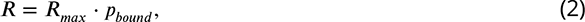

where *R_max_* is a constant coefficient that dictates the maximum possible polymerase loading rate. Thermodynamic models enable the calculation of *p_bound_* by assigning a statistical weight to each possible state in which the regulatory system can be found. In the case of a minimal enhancer with one activator binding site, the enhancer-promoter DNA can be empty, occupied by Dorsal, occupied by RNAP, or simultaneously bound by Dorsal and RNAP (Fig. 2B). The statistical weight associated with each of these terms is shown in Figure 2B. Here, [*Dl*]/[*K_D_* ] is the statistical weight associated with 1nding Dorsal (with concentration [*Dl*] and binding dissociation constant *K_D_* ) bound to the promoter alone, while [*P* ]/[*K_P_* ] is the weight of finding RNAP (with concentration [*P* ] and binding dissociation constant *K_P_* ) bound to the promoter alone. Note that the weight of having both Dorsal and RNAP bound simultaneously includes an extra glue-like cooperativity coefficient, *w*, that determines how strongly Dorsal recruits RNAP to the promoter. The value of *ω* is constrained to be > 1 so that higher Dorsal occupancy leads to higher RNAP occupancy.

To calculate *p_bound_* , we divide the sum of the weights featuring a bound RNAP molecule by the sum of all possible weights. Substituting this calculation into Equation 2 yields

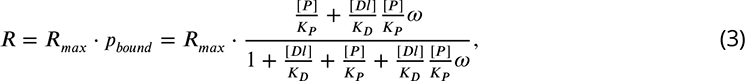

which is plotted in Figure 2E. As shown in the 1gure, increasing *K_D_* shifts the concentration at which the RNAP loading rate reaches half its maximum value toward higher Dorsal concentrations, but does not change the overall shape of the curve. We also note the presence of a non-zero baseline of RNAP loading rate due to the Dorsal-independent [*P* ]/[*K_P_* ] term in the numerator of Equation 3. This baseline suggests that it could be possible for a promoter in the ’ON’ state to produce low, basal-level transcription in the absence of bound Dorsal.

Together, the kinetic barrier model outlined in Figure 2A and the thermodynamic model’s Equation 3 de1ne a comprehensive quantitative framework that predicts how the fraction of active loci, the transcriptional onset time, and the RNAP loading rate as a function of Dorsal concentration vary as model parameters such as the Dorsal dissociation constant *K_D_* are modulated (Fig. 2D,E). These predictions constitute hypotheses that we experimentally tested throughout the remainder of this work.

### 2.2 Establishing a minimal synthetic enhancer system to test theoretical predic- tions

To test our model’s predictions, we constructed single binding site enhancers driven by the Dorsal activator. Dorsal is one of the best characterized transcription factors in *Drosophila* and a classic example of a morphogen (***Roth et al., 1989***; ***Reeves et al., 2012***). Dorsal is provided maternally and forms a dorsoventral gradient of nuclear localization (Fig. 3A) (***Gilbert, 2010***), acting as an activator by default (***Thisse et al., 1991***; ***Jiang et al., 1991***) and as a repressor in the presence of nearby binding sites for corepressors (***Kirov et al., 1993***; ***Papagianni et al., 2018***). Prior to activation of the zygotic genome (up to the 12th mitotic cycle), Dorsal is the only transcription factor with a nuclear protein gradient across the dorsoventral axis (***Sandler and Stathopoulos, 2016***; ***Dufourt et al., 2020***). Thus, the Dorsal nuclear concentration is the sole source of dorsoventral positional information for developmental enhancers at this stage in development. These features, combined, make Dorsal an ideal input transcription factor for activating a minimal synthetic reporter system.

**Figure 3.**
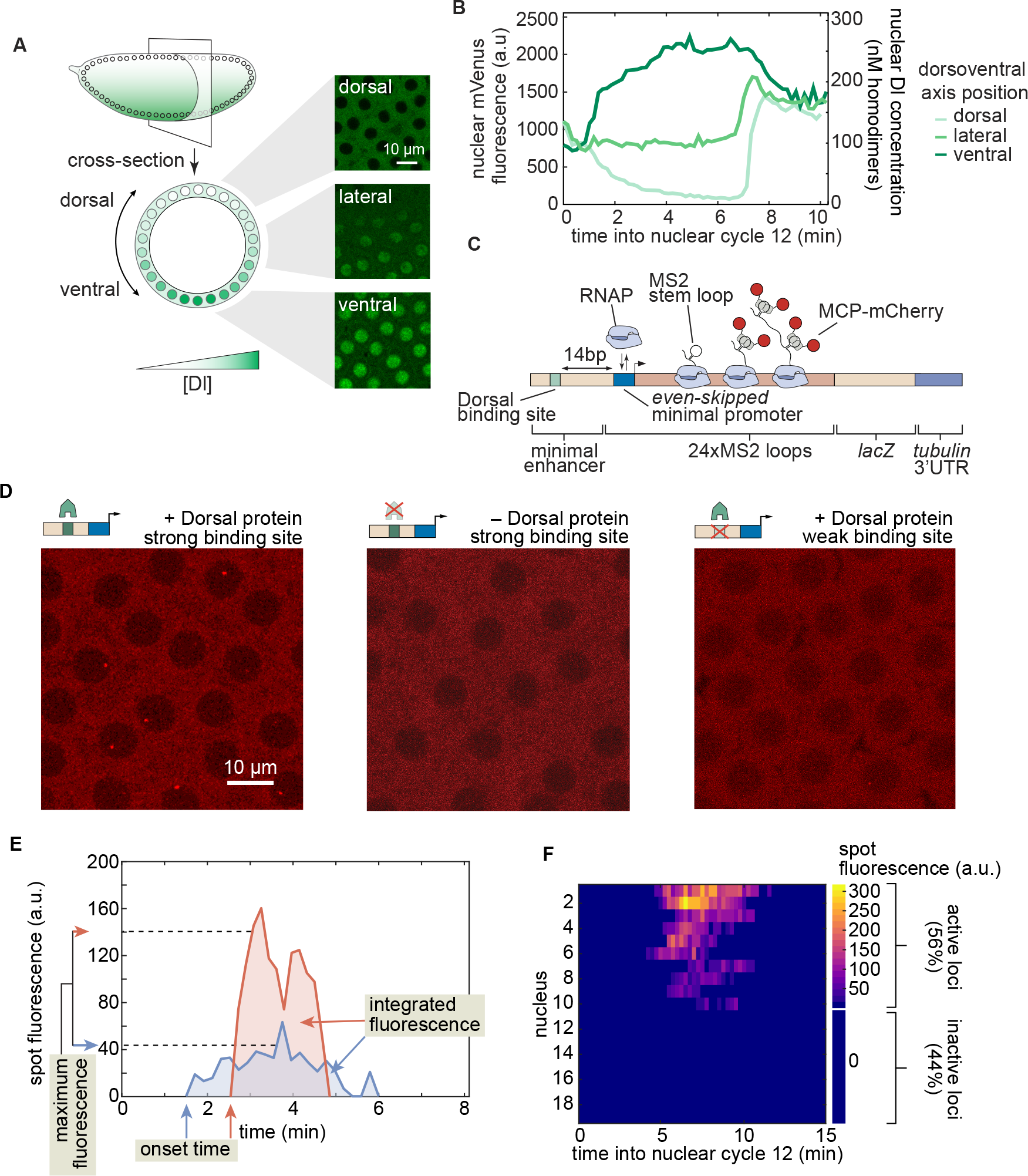
Simultaneously measuring transcription factor protein input and transcriptional output. **(A)** Schematic of the Dorsal protein gradient in early *Drosophila* embryos. Dorsal protein accumulates in ventral nuclei and is progressively excluded from more dorsal nuclei. Example snapshots show Dorsal-mVenus in various positions along the dorsoventral axis. **(B)** Representative time traces of nuclear Dorsal-mVenus 2uorescence in various positions along the dorsoventral axis. The right y-axis shows the nuclear Dorsal concentration according to the calibration described in Figure S6. **(C)** Schematic of minimal synthetic enhancer system containing a single binding site for Dorsal that drives transcription of a reporter tagged with MS2 loops, which are visualized through the binding of MCP-mCherry. The Dorsal binding site is placed 14 bp upstream of the *even-skipped* minimal promoter. **(D)** Snapshots from embryos containing an optimal binding-site reporter in the presence (left) or absence (middle) of Dorsal, or containing a strongly mutated Dorsal binding site (right). **(E)** Example 2uorescence time traces and quantitative metrics of transcriptional activity. **(F)** Fluorescence of all transcription spots in individual nuclei in the 1eld of view of one embryo as a function of time. If a transcription spot was detected within a nucleus at any point during the interphase of nuclear cycle 12, then the locus was considered active; otherwise, the locus was classi1ed as inactive.

In order to relate output transcriptional activity to the time-variant input Dorsal concentration throughout development, we measured the instantaneous Dorsal concentration in live embryos by creating a CRISPR knock-in Dorsal-mVenus fusion allele based on a previous Dorsal fusion (***Reeves et al., 2012***) that rescues embryonic development (***Kremers et al.*** (***2006***); ***Gratz et al.*** (***2015***); Materials and methods). Further, in order to increase the dynamic range of Dorsal concentration in our experiments, we further combined this CRISPR allele with a Dorsal-mVenus transgene (***Reeves et al., 2012***), resulting in a line that will hereafter be referred to as 2x Dorsal flies. This fusion made it possible to quantify the concentration dynamics of the Dorsal protein input (Fig. 3A,B) in individual nuclei (Video S4 , left; Materials and methods). Dorsal-mVenus nuclear 2uorescence time traces quanti1ed over nuclear cycle 12 confirmed the dynamic nature of Dorsal concentration and were quantitatively similar to previous measurements (Fig. 3B; ***Reeves et al.*** (***2012***); details of Dorsal-mVenus quantification in Fig. S5A,B). Nuclear cycle 12 nuclei in 2x Dorsal 2ies experience a Dorsal concentration gradient spanning multiple orders of magnitude, from less than 1 nM to ≈ 400 nM (Fig. 3B; details of Dorsal-mVenus calibration in Fig. S6).

To visualize the dynamics of Dorsal-dependent transcription, we developed a reporter transgene containing a minimal synthetic enhancer consisting of a single high affinity, consensus binding site for the Dorsal transcription factor (***Ip et al., 1992***; ***Jiang and Levine, 1993***; ***Szymanski and Levine, 1995***) (Fig. 3C). Hereafter we refer to this strong site enhancer as as DBS_6.23 for <u>D</u>orsal <u>B</u>inding <u>S</u>ite, followed by its binding affinity score according to the Patser algorithm (***Stormo and Hartzell*** (***1989***); Materials and methods). To quantify the transcriptional activity of this enhancer, we used the MS2-MCP system to fluorescently label nascent RNA molecules in our reporter constructs, which appear as nuclear fluorescent puncta (hereafter “transcription spots”) in laser-scanning confocal microscopy movies (Video S4 , right; ***Bertrand et al.*** (***1998***); ***Garcia et al.*** (***2013***); ***Lucas et al.*** (***2013***). We performed image analysis of the MS2 movies using a custom data analysis pipeline in Matlab and Fiji (Materials and methods; (***Schindelin et al., 2012***; ***Lammers et al., 2020***).

To validate this minimal synthetic system, we determined that DBS_6.23-MS2 drives detectable and quanti1able levels of transcription, and that this transcriptional activity is mainly governed by Dorsal. We compared the transcriptional activity of DBS_6.23-MS2 in embryos laid by 2x Dorsal females with the activity in embryos laid by females homozygous for a *dorsal* null allele. While transcription spots were clearly present in the 2x Dorsal background (Fig. 3D, left), they were extremely rare in *dorsal* null embryos (Fig. 3D, middle): not a single transcription spot was detected during nuclear cycle 12 in any of 4 replicates containing *>* 60 nuclei in total. Dorsal is therefore necessary for transcriptional activity in our reporter constructs.

We next sought to determine whether the detected transcriptional activation is solely due to Dorsal interacting with the binding site explicitly engineered into the construct or whether there are cryptic Dorsal binding sites contributing to gene expression. We generated a second reporter, DBS_4.29-MS2 in which the Dorsal binding site was strongly perturbed using known point mutations (***Ip et al., 1992***). Transcription was rarely detectable in DBS_4.29-MS2 embryos (Fig. 3D, right), with the average transcriptional activity (mean instantaneous fluorescence) per detected spot being less than 10% of the optimal DBS_6.23 enhancer at any Dorsal concentration (Fig. S9). Thus, the Dorsal site placed within the synthetic enhancer is necessary for robust activation and is the main driver of this transcriptional activity.

Next, we identified which observable features in the MS2 signal could be used as metrics for quantifying Dorsal-dependent transcriptional activity. We collected DBS_6.23-MS2 time traces of MCP-mCherry fluorescence from transcription spots during nuclear cycle 12 along with four metrics of transcriptional activity (Fig. 3E,F). First, the maximum spot fluorescence corresponds to the 95th percentile of intensity over time, which is proportional to the transcription rate (Section S1 .2). Second, the transcriptional onset time is defined as the time since the previous mitosis at which a transcription spot is first detected (Fig. S3). Third, the integrated spot fluorescence corresponds to the time integral of the spot fluorescence and is directly proportional to the amount of mRNA produced by the locus (***Garcia et al., 2013***) (Materials and methods). Finally, as previously observed in other genes in flies (***Garcia et al., 2013***; ***Dufourt et al., 2018***; ***Lammers et al., 2020***; ***Harden et al., 2021***), not all nuclei exposed to the same average nuclear Dorsal concentration exhibited detectable transcription (Fig. 3F). As a result, we quantified the fraction of active loci—regardless of their level of activity or temporal dynamics—by measuring the number of nuclei with observable transcription signal in at least one movie frame throughout nuclear cycle 12, divided by the total number of nuclei in the field of view. Thus, we have established quantitative metrics that enable us to engage in a dialogue between experiment and a theory of Dorsal-driven transcriptional dynamics.

### 2.3 Transcriptionally active and inactive loci correspond to functionally distinct populations

Before attempting to predict Dorsal-driven transcriptional dynamics, it is important to ensure that the fact that only some loci engage in transcription is the result of Dorsal action and not of limitations of our experimental setup. Transcriptionally silent loci that remain inactive throughout interphase, such as those revealed by our experiment (Fig. 3F), have been observed using MS2 (and its sister mRNA labeling tool, PP7) in live-imaging experiments in flies (***Garcia et al., 2013***; ***Lammers et al., 2020***; ***Berrocal et al., 2020***), plants (***Alamos et al., 2020***), and mammalian cells (***Hafner et al., 2020***). However, it has not been possible to determine whether these inactive loci correspond to a separate transcriptional state from active loci, or whether they are an artifact of the fluorescence detection thresholds associated with various microscopy techniques.

To answer this question, it is necessary to quantify MS2 fluorescence at these inactive loci and determine whether they differ from loci not exposed to activators, which do not transcribe (Fig. 3F). However, to date this approach has not been feasible because most MS2 measurements have relied on the presence of an MS2 signal itself to segment and quantify the fluorescence of transcription spots. We hypothesized that, if undetected loci correspond to a distinct and weaker, Dorsal-independent state, then detected and undetected spots in embryos carrying wild-type Dorsal would appear as two distinct populations. In this scenario, the mCherry fluorescence of undetected spots corresponding to inactive loci in wild-type Dorsal embryos would be similar to that observed in Dorsal null embryos, and clearly distinct from the mCherry fluorescence of active loci in the presence of Dorsal.

To quantify MS2 fluorescence independently of whether a MS2 spot was detected, we implemented the *parS*-ParB DNA labeling system (***Germier et al., 2017***; ***Chen et al., 2018***). Here, fluorescently labeled ParB proteins bind the parS DNA sequence resulting in a fluorescence spot appearing at the locus independently of the transcriptional state of the locus (Fig. 4A). We created flies with and without functional Dorsal expressing ParB2-eGFP (subsequently referred to as ParB-eGFP) and MCP-mCherry to label our locus DNA and nascent RNA, respectively. We crossed flies containing parS-DBS_6.23-MS2 to flies carrying ParB-eGFP and MCP-mCherry to generate embryos that have our locus of interest labeled with ParB-eGFP colocalized with the transcriptional signal in the MCP-mCherry channel (Fig. 4A,B; Video S4 ).

**Figure 4.**
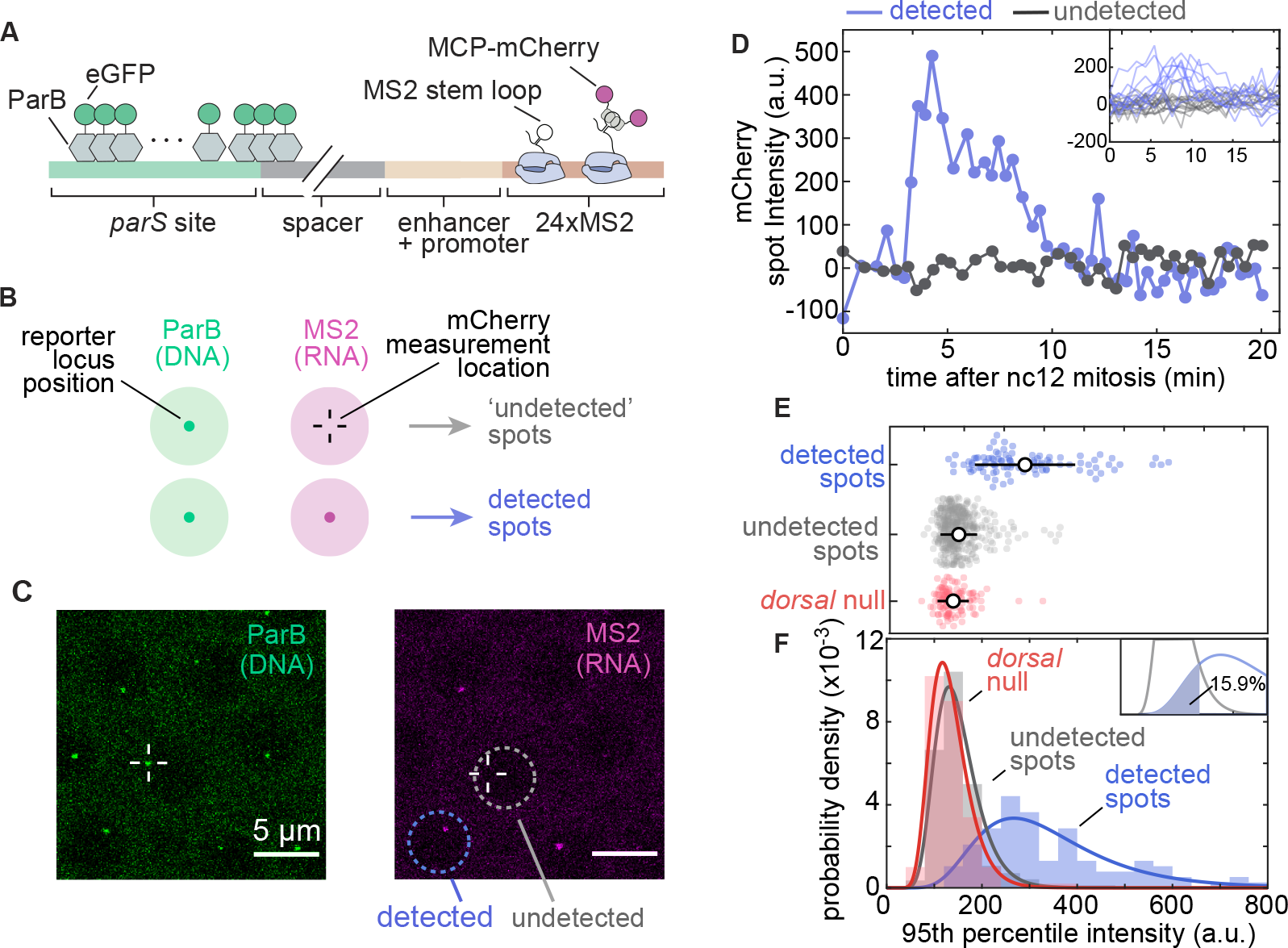
Transcriptionally independent ParB labeling con1rms that transcriptionally inactive loci are functionally distinct from active loci. **(A)** Schematic of ParB-eGFP construct. ParB-eGFP molecules bind and polymerize out from *parS* sequences, which are placed "’ 400 bp upstream of the enhancer. The enhancer and promoter together drive transcription of MS2 loops that subsequently bind MCP-mCherry. **(B)** Schematic of the experiment. Loci are located by detecting a signal in the ParB-eGFP channel; these locations were used to 1t a 2D Gaussian to the same area in the MS2-mCherry channel to estimate 2uorescence intensity regardless of whether an MS2-mCherry signal was detected (Materials and methods Sec. 4.4). **(C)** Example images of ParB-eGFP (left) and MCP-mCherry (right) channels. Detected and undetected transcriptionally active loci solely based on the MCP-mCherry signal alone are shown. **(D)** Example time traces of MCP-mCherry 2uorescence over time at the ParB-eGFP loci in nuclei with (blue) and without (grey) detected MS2-mCherry spots of the DBS_6.23 enhancer showing clear qualitative differences between the two populations. Inset, all detected and undetected 2uorescence traces obtained in the same embryo. Negative intensity values are due to spot intensities very close to the background 2uorescence. **(E)** Swarm plots of 95th percentile MCP-mCherry 2uorescence at loci with detected (blue; N = 125) and undetected MS2-mCherry transcription (gray; N = 425) driven by the DBS_6.23 enhancer in wild-type Dorsal embryos. Red (N = 96), maximum 2uorescence of all loci in Dorsal null embryos, de1ned as the 95th percentile of intensity over time (black circles, mean; bars, standard deviation). Detected spots are signi1cantly different from both null (ANOVA, p < 0.01) and undetected spots (ANOVA, p < 0.01) **(F)** Histograms of the data shown in (E). Solid lines correspond to log-normal 1ts performed for ease of visualization. Inset, undetected and detected distribution 1ts and the area used to estimate the false-negative detection rate of 15.9%.).

Guided by the spatial positions reported by ParB-eGFP, we measured the MCP-mCherry signal at all DBS_6.23 reporter loci in embryos carrying wild-type Dorsal (Fig. 4C) or laid by mothers homozygous for the *dl*^1^ null allele (Dorsal null embryos). We then classified loci from wild-type Dorsal embryos into two categories, detected and undetected, depending on whether they were identified as spots in the MCP-mCherry channel by our analysis pipeline (Fig. 4B,C; Section 4.5). As shown in the the examples presented in Figure 4D, there are clear qualitative differences between MCP-mCherry fluorescence time traces corresponding to detected or undetected transcriptional spots from wild-type embryos. Thus, our analysis made it possible to quantify MS2 fluorescence in three populations: all loci in Dorsal null embryos, undetected loci in wild-type Dorsal embryos, and detected loci in wild-type Dorsal embryos.

To compare these populations, we computed the 95th percentile value over each locus’ MCP- mCherry 2uorescence time trace (Fig. 4E). The distribution of mCherry fluorescence from undetected spots in wild-type Dorsal embryos largely overlapped with that of all spots in Dorsal-null embryos (Fig. 4F), consistent with these two populations corresponding to loci expressing Dorsal-independent levels of activity. Moreover, both distributions were clearly distinct from the distribution of detected spots in wild-type Dorsal embryos (Fig. 4E,F). Thus, our results provide strong evidence that inactive loci are not artifacts of the detection limit of our imaging techniques. Rather, loci can belong to one of two distinct populations: those that transcribe at a high, Dorsal-dependent level and those that are transcriptionally inactive (or active at a low, undetectable level that is comparable to that of embryos lacking Dorsal). We therefore conclude that the decision to transcribe made by each locus is an additional regulatory strategy controlled by Dorsal.

From the observations in Figure 4E and F, we estimated our error in classifying loci as inactive. This false-negative detection rate, corresponding to the area under the curve shaded in the inset of Figure 4F, is estimated as 15.9%. However, this false-negative rate is likely an underestimation. For example, this rate may depend on Dorsal concentration, which cannot be controlled for in this experiment. Additionally, the presence of ParB in the locus may itself affect transcriptional dynamics, impacting the false-negative rate. For these reasons, we do not attempt to correct our measurements of the fraction of active loci using this estimated false-negative rate.

### 2.4 Dorsal-dependent kinetic barriers explain transcription onset dynamics and modulation of the fraction of active loci

Having established that transcriptionally inactive promoters mostly constitute a separate population from transcriptionally active promoters (Fig. 4), we sought to test whether our theoretical model (Fig. 2A) can quantitatively recapitulate the fraction of active loci and their transcription onset times. Tuning transcription factor-DNA binding affinity has been a powerful tool to test models of transcriptional regulation in the past (***Meijsing et al., 2009***; ***Phillips et al., 2019***). Inspired by these previous works, we probed our model by adjusting the Dorsal-DNA interaction energy in our minimal synthetic enhancer.

We constructed a series of enhancers containing a single binding site with varying affinities for Dorsal. Building on the optimal DBS_6.23 and the mutated DBS_4.29 sites (Fig. 3D, left vs. right), we created 1ve additional enhancers of varying intermediate strengths by introducing point mutations into the consensus Dorsal binding motif to obtain a range of predicted affinities (Fig. 5A,B; Materials and methods Section 4.1). As described above, we refer to these enhancers as DBS, followed by their corresponding Patser score.

**Figure 5.**
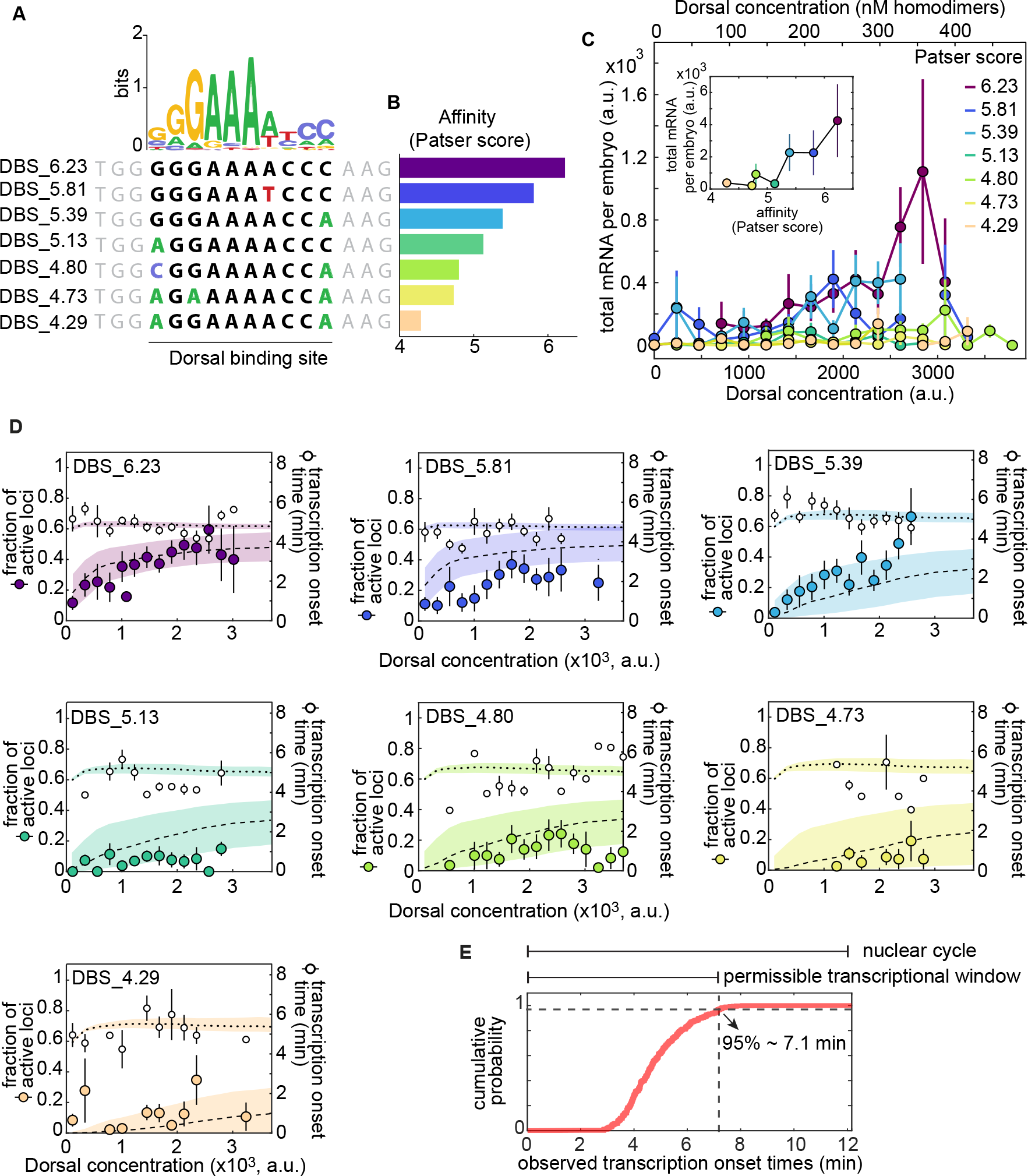
A multi-step kinetic barrier model predicts the Dorsal-dependent fraction of active loci with constant mean transcriptional onset times. **(A)** Top: Dorsal positional weight matrix logo from *Ivan et al.* (*2008*). Bottom: Sequence of the Dorsal binding sites engineered into minimal synthetic enhancers. Bold letters, 10 bp Dorsal motif. Black letters, consensus bases; colored letters, mutated bases; gray letters, sequence context. **(B)** Relative affinities of Dorsal binding sites estimated from the Patser algorithm using the Dorsal position weight matrix. **(C)** Overall transcriptional activity driven by the enhancers containing the binding sites in (A) measured as the total produced mRNA (2uorescence integrated over nuclear cycle 12) as a function of Dorsal concentration. Inset, mean total mRNA produced per embryo integrated across all Dorsal concentrations. Error bars, SEM over N > 3 embryos containing 3 or more nuclei belonging to that 2uorescence bin. The top x-axis shows the estimated nuclear Dorsal concentration according to the calibration described in Figure S6. **A multi-step kinetic barrier model explains the Dorsal-dependent fraction of active loci with constant mean transcriptional onset times. (D**) Data and model 1ts for the fraction of active loci (left y-axis) and mean transcription onset time (right y-axis) for each enhancer. Empty black circles, experimentally observed mean transcription onset time; 1lled circles, experimentally observed mean fraction of active loci. Fitted curves are represented as dashed lines (fraction of active loci) and dotted lines (mean onset times), corresponding to predictions using median parameter values from the joint posterior distribution. Shaded areas, 95% credible interval (see Table S1 for inferred parameter values). Error bars, SEM over N > 3 embryos containing 3 or more nuclei belonging to that 2uorescence bin. **(E)** Cumulative distribution of mean spot detection times per Dorsal 2uorescence bin across all embryos and enhancers (N = 344 spots). Vertical dashed line, time at which 95% of spots have turned on (≈ 7.1 min) and end of the permissible transcription time window.

For the purpose of quantifying output transcriptional activity as a function of Dorsal concentration, we assigned a single Dorsal concentration value to each nucleus corresponding to the mVenus fluorescence in the center of that nucleus at a fiducial time point halfway through each nucleus’s lifetime, approximately in the middle of nuclear cycle 12 when Dorsal levels are relatively stable (Fig. S5A,B). We next grouped nuclei into 17 linearly spaced bins that span the dorsoventral axis based on their 1ducial 2uorescence (Fig. S5B).

We assessed whether these point mutations were sufficient to generate a graded response to Dorsal and to determine the dynamic range of gene expression afforded by these enhancers. To make this possible, we integrated the total mRNA output over nuclear cycle 12 of each enhancer as a function of Dorsal concentration across all nuclei exposed to a given Dorsal concentration. The integrated mRNA output of the four weakest enhancers changed little across the dorsoventral axis (Fig. 5C). However, an appreciable trend in integrated mRNA was observed for the three strongest affinities (Fig. 5C). Further, plotting the total mRNA integrated across the entire dorsoventral axis of the embryo as a function of Patser score revealed that binding-site affinity (as reported by Patser score) is strongly correlated with transcriptional output in our single binding site enhancers (Fig. 5C, inset). In the case of this measure, there was also a threshold affinity: enhancers containing binding sites with affinities below that of DBS_5.13 showed no substantial differences in transcriptional activity (inset, Fig. 5C).

We used these constructs to measure mean transcriptional onset time as a function of Dorsal concentration and binding affinity, one of the key magnitudes predicted by our model (Fig. 2D). The measured mean onset time was relatively constant at ≈5 minutes across all Dorsal concentrations and enhancer constructs (Fig. 5D, dotted lines). This value is consistent with the measured onset times of other early embryonic genes such as the minimal *hunchback* promoter P2P (***Garcia et al., 2013***; ***Lucas et al., 2013***; ***Eck et al., 2020***).

We also determined that the fraction of active loci is highly sensitive to Dorsal concentrations and Dorsal binding-site affinity (Fig. 5D, dashed lines). The strongest Dorsal binding sites showed a large modulation of the fraction of active loci across Dorsal concentrations, while the weakest drove a relatively constant and low fraction of active loci across all Dorsal concentrations (Fig. 5D). Our kinetic barrier model assumes that loci which fail to become active during the permissible transcription time window will remain inactive during the rest of the nuclear cycle (Fig. 2C). As a result, to determine whether the kinetic barrier model recapitulates the observations in Figure 5D, it was necessary to assign a value to this time window. We reasoned that the end of this time window determines the time point at which new transcription spots can no longer appear, possibly due to the onset of the next round of mitosis. To estimate the time point when nearly all spots have turned on, we calculated the 95th percentile of the observed spot onset times across all affinities: ≈ 7.1 min after the previous anaphase (Fig. 5E).

Using the measured time window of permissible transcription, we performed a simultaneous 1t to the fraction of active loci and mean transcription onset times across all enhancers using the kinetic barrier model from Section 2.1 (Fig. 5D). Consistent with our model, we forced all enhancers to share the same value for *c*, and only letting the Dorsal dissociation constant, *K_D_* , vary for each enhancer separately. By systematically exploring models with different numbers of OFF states *n* (Fig. S10, Fig. S11), we determined that a biochemical cascade with at least 3 to 4 rate-limiting OFF states is capable of capturing the qualitative behavior of our observations: a Dorsal concentration- and binding affinity-dependent fraction of active loci (dashed lines in Fig. 5D) and a mean transcription onset time that is mostly constant across Dorsal concentrations and affinities (dotted lines in Fig. 5D). Interestingly, alternative functional forms for *k*, such as modeling this transition rate as depending linearly on Dorsal concentration, instead of depending on Dorsal DNA occupancy, resulted in worse 1ts to the fraction of active loci at saturating concentrations of Dorsal (Section S1 .5; Fig. S4). Thus, our observations can be explained by a model in which Dorsal, through DNA binding, accelerates the promoter’s transition through a sequence of kinetic barriers to a state of active transcription.

### 2.5 The experimentally measured RNAP loading rate are compatible with a thermodynamic binding model

As a next step in our theoretical dissection, we tested the performance of our theoretical model in explaining the rate of transcription after loci become active. Typically, in MS2 experiments, the loading rate is measured from the initial slope of spot fluorescence traces (***Garcia et al., 2013***; ***Eck et al., 2020***; ***Liu et al., 2021***). However, due to the weak expression driven by our enhancers, it was not possible to perform this analysis with confidence (Fig. S8). In lieu of directly measuring the transcription rate, we evaluated a related, more robust and readily observable quantity: the maximum trace fluorescence (Fig. 3E). We approximately relate the RNAP loading rate predicted by the simple activator model (Equation 3) to the maximum fluorescence by a constant factor (Appendix S1 .2), enabling direct comparison between theoretical predictions and experimental data.

Measurements of the maximum spot fluorescence over time as a function of Dorsal concentration for each of our seven minimal synthetic enhancers revealed that the maximum fluorescence is relatively constant across Dorsal concentration for most binding sites—particularly for the weakest of them, DBS_5.13, DBS_4.73, and DBS_4.23 (Fig. 6). However, the sparse and noisy nature of our data makes it challenging to draw confident conclusions from the 1ts, even for the stronger binding sites (i.e. DBS_6.23, DBS_5.81, and DBS_5.39). In the case of the lower affinity binding sites, the constant maximum fluorescence suggests that the Dorsal concentration level in our embryos is far below the Dorsal dissociation constant *K_D_* , even after increasing the Dorsal dosage by a factor of two as in our 2x Dorsal line. The effect of very low Dorsal concentrations relative to their respective *K_D_* values can be clearly seen in Equation 3 and in Figure 2, where, for [*Dl*]/*K_D_ «* 1, the RNAP loading rate, *R*, adopts a basal level given by

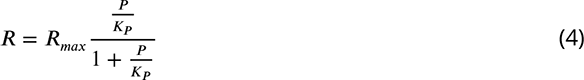

that is independent of Dorsal concentration and binding affinity.

**Figure 6.**
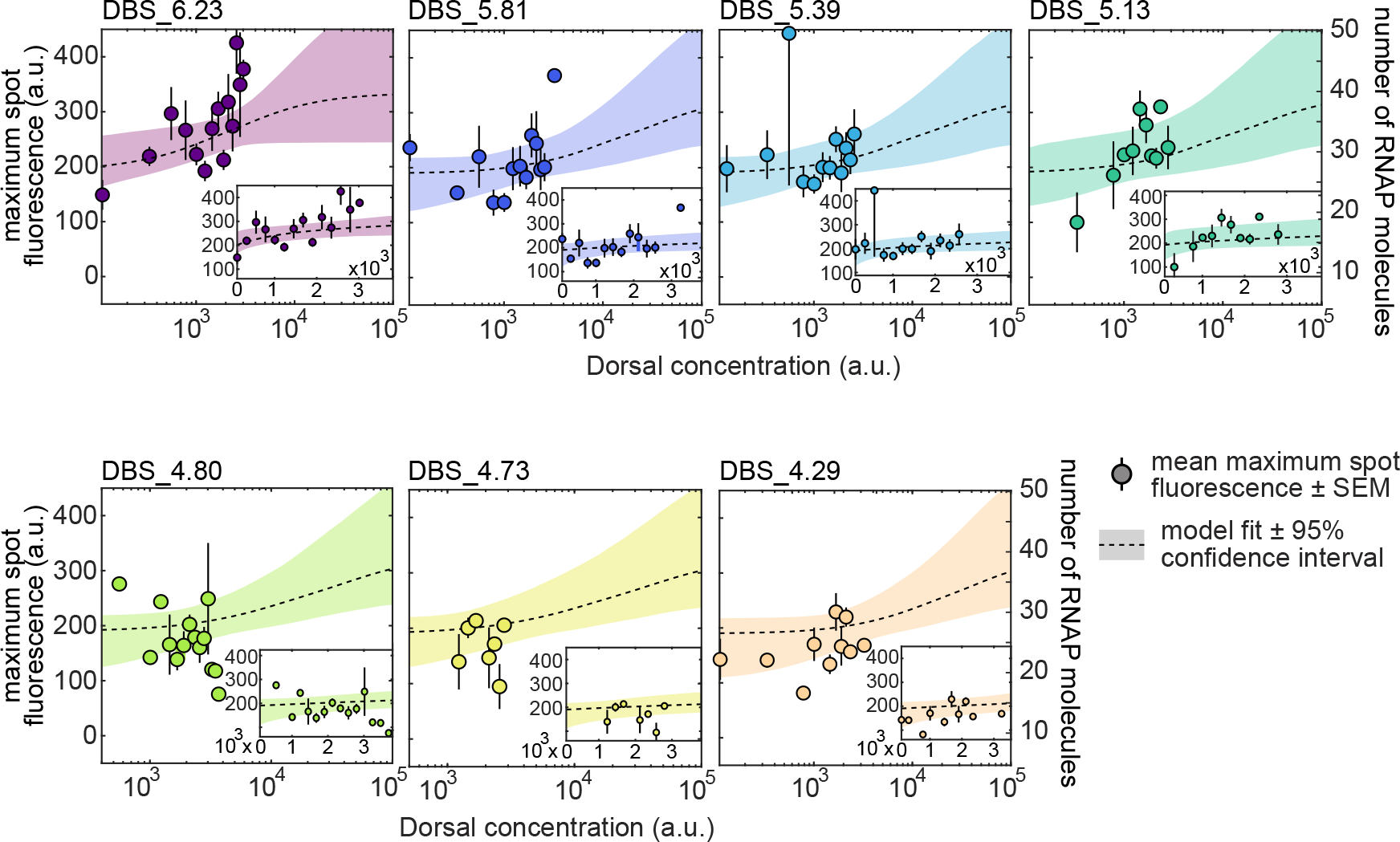
Testing RNAP loading rate predictions of the thermodynamic model. Mean maximum spot 2uorescence as a function of Dorsal concentration for minimal synthetic enhancers with different affinities for Dorsal (1lled circles). The right y-axis denotes the calibrated number of actively transcribing RNAP molecules (for details of calibration, see Section S1 .3 and Fig. S2). Dashed curves correspond to a simultaneous Markov Chain Monte Carlo curve 1t to all data using Equation 3. Fits share all parameters except *K_D_* . Shaded areas, 95% prediction intervals. Insets, same data and 1ts plotted on a linear scale with axis ranges zoomed in on the data. See Table S2 for inferred parameter values. Error bars, SEM across N > 3 embryos containing 3 or more nuclei in a given 2uorescence bin.

As shown on the right y-axes in Figure 6, this basal level corresponds to ≈ 20 RNAP molecules actively transcribing the gene (≈ 15% of the maximum number of RNAPs that can 1t on the gene, as described in Section S1 .3). For ease of visual comparison to the thermodynamic model predictions, we also plotted best-1t theoretical curves on top of the data using dashed curves (the insets in Fig. 6 show the same plots but zoomed into the measured data and plotted on a linear scale). These 1ts further underscore that our data do not explore a wide dynamic range with the precision necessary to determine the magnitude of *K_D_* for each construct and to thoroughly test the thermodynamic model.

## 3 Discussion

A major obstacle to uncovering the mechanistic and quantitative underpinnings of enhancer action is the inherent complexity of endogenous regulatory sequences. Synthetic minimal enhancers are powerful alternatives to the complex experimental reality faced by modeling efforts in endogenous enhancers (***Garcia et al., 2016***, ***2020***). Synthetic minimal enhancers contain binding sites for one or a handful of transcription factors, making them more amenable to theoretical dissection (***Fakhouri et al., 2010***; ***Sayal et al., 2016***; ***Crocker and Ilsley, 2017***) and revealing the complex interplay among activators, repressors, and pioneer factors, as well as their contribution to mRNA transcript accumulation (***Fakhouri et al., 2010***; ***Sayal et al., 2016***; ***Crocker and Ilsley, 2017***). However, previous synthetic-based efforts to dissect enhancer function always involved fixed-embryo measurements, which cannot reveal the three inherently dynamical roles dictated by enhancer sequences (Fig. 1). Here we augmented previous synthetic approaches by quantifying the real-time action of minimal enhancers with one binding site for the Dorsal activator in single cells of living, developing *Drosophila* embryos using the MS2 system. Contrary to theoretical speculations that single binding sites within eukaryotic genomes lack enough information to be recognized by transcription factors in the absence of other nearby binding sites (***Wunderlich and Mirny, 2009***), we demonstrated that Dorsal can drive expression when bound to single binding sites (Fig. 3D). Additionally, we demonstrated that the fraction of active loci is a feature under regulatory control in our synthetic system (Fig. 3F; Fig. 4F), confirming the important role of this regulatory strategy in shaping the expression dynamics of endogenous enhancers (***Garcia et al., 2013***; ***Dufourt et al., 2018***; ***Lammers et al., 2020***; ***Harden et al., 2021***). Thus, while the signal driven by our minimal synthetic constructs is weak (Fig. 6), it can be quantified and recapitulates biologically relevant dynamic features of transcription that are also at play in endogenous enhancers.

It is important to note that the uncovering of a fraction of inactive loci in many reporter systems by us and others (***Garcia et al., 2013***; ***Dufourt et al., 2018***; ***Lammers et al., 2020***; ***Harden et al., 2021***) did not necessarily imply that this modulation of transcriptional engagement constitutes a biological control variable. Indeed, because live cell imaging techniques typically lack single-molecule resolution, it was unclear whether undetected loci in our study—and all previous studies— corresponded to a distinct population or were a detection artifact. By simultaneously labeling the locus with the transcription-independent reporter ParB-eGFP and nascent mRNA with MCP-mCherry (Fig. 4A), we demonstrated that a significant number of loci categorized as inactive do not constitute an experimental artifact and instead correspond to a distinct transcriptional state that is comparable to that measured in the absence of Dorsal protein (Fig. 4). In the future, conducting all live transcription measurements with DNA loci labeled by ParB could make it possible to confidently quantify the activity of all loci regardless of their activity.

Our minimal synthetic constructs and our validation of a distinct population of inactive loci enabled us to test an emerging theoretical model of enhancer action in development: a kinetic barrier model of transcriptional engagement (Fig. 2A; ***Fritzsch et al.*** (***2018***); ***Dufourt et al.*** (***2018***); ***Eck et al.*** (***2020***)). Importantly, our model deviated from previous theoretical efforts that assumed that the transition rates between states preceding transcriptional engagement were either constant (***Dufourt et al., 2018***) or depended linearly on activator concentration (***Eck et al., 2020***). Instead, in order to account for the effects of Dorsal binding affinity on transcriptional dynamics, we assumed that this rate was proportional to Dorsal occupancy at its target DNA site. Thus, while the mechanisms underlying several aspects of this model, such as the molecular identity of the various OFF states, remain unknown, this model can generate predictions for how the fraction of active loci and the transcriptional onset time are modulated by the Dorsal concentration and its binding affinity (Fig. 2C-E).

We systematically challenged this model by generating a small collection of minimal synthetic enhancers spanning a large range of affinities for Dorsal (Fig. 5A). Comparing the fraction of active loci and the transcription onset times of these enhancers revealed that the kinetic barrier model recapitulated our measurements (Fig. 5D). In past studies probing transcription dynamics in the *Drosophila* embryo (***Dufourt et al., 2018***; ***Eck et al., 2020***), the pioneer factor Zelda was found to be largely responsible for ensuring constant transcription factor onset times and for determining the fraction of active loci. We cannot rule out the potential existence of distant or low-affinity Zelda binding sites (***Rushlow and Shvartsman, 2012***) in our constructs. Alternatively, as it was recently demonstrated for the Bicoid activator ***Hannon et al.*** (***2017***), Dorsal could also have a pioneering activity. Indeed, the Dorsal homolog NF-*,c*B has been recently shown to displace nucleosomes (***Cheng et al., 2021***). To further test the kinetic barrier model, it would be informative to directly perturb the temporal dynamics of nuclear Dorsal concentration to affect transcriptional engagement. For example, several optogenetics systems have been successfully deployed in the early 2y embryo to inactivate transcription factors during discrete time widows (***Huang et al., 2017***; ***McDaniel et al., 2019***; ***Irizarry et al., 2020***). In the future, a version of one of these systems may dissect how the temporal dynamics of Dorsal concentration affect transcriptional activation.

Although the kinetic barrier model predicted the fraction of active loci and onset times (Fig. 5D) relatively well, we were unable to use our data to conclusively test the thermodynamic model’s predictions of the rate of mRNA production (Fig. 6). Such limitation stemmed from the fact that only a fraction of loci display detectable transcription that can be used to quantify the mRNA production rate. Further, among these loci, the rate of transcription was found to be highly variable. As a result, our statistics were limited such that it was not possible to perform an unequivocal test of the thermodynamic model.

The apparent lack of substantial Dorsal concentration dependence observed in our measurements of RNAP loading rate could be explained in two possible ways. First, it is possible that there is a modulation of this rate in our measurements, but that this modulation is obscured by our experimental noise. Second, the Dorsal concentrations accessed by our experiment could be below the *K_D_* of our binding sites. In this scenario, a modulation in the mRNA production rate would become apparent only at Dorsal concentrations higher than those attainable by our experimental system. While our embryos contained double the genetic dosage of Dorsal compared to wild type, perhaps 5-10 times the wild-type Dorsal concentration could be needed to exceed the *K_D_* and modulate the rate of mRNA production. To express this high Dorsal concentration, which is certain to affect normal embryonic development, genetic approaches to increase Dorsal dosage in the embryos similar to those recently applied to flatten the Bicoid gradient might be necessary (***Hannon et al., 2017***).

It is important to note that, despite not seeing a modulation in the rate of mRNA production, we do see a significant change in the fraction of active loci as Dorsal concentration is varied (Fig. 5). This seeming contradiction could be explained through the presence of two dissociation constants in our model (Fig. 2): one dissociation constant for the first part of the model governing the onset of transcription, and a different dissociation constant for the second part of the model dictating the rate of RNAP loading once transcription has ensued. Interestingly, previous works quantifying transcriptional dynamics of a minimal Bicoid-activated *hunchback* P2 enhancers also hint at the existence of these two distinct dissociation constants (***Garcia et al., 2013***).

Further, this model is consistent with our surprising observation of a basal level of transcription in the presence of even extremely weak binding sites (Fig. 6) despite the lack of detected transcription in the absence of Dorsal protein (Fig. 3D, middle). This observation could be explained if Dorsal acted as both as a pioneer-like transcription factor triggering the onset of transcription, even at low concentrations relative to its *K_D_* , and as an activator of the transcription rate at high concentrations. Going forward, synthetic minimal enhancers could constitute the foundation for exploring the behavior of more complex regulatory regions. Independently inferring biophysical parameters such as Dorsal-DNA binding and dissociation constants could help constrain models of Dorsal participating in the activation of promoters with additional activators and repressors (***Fakhouri et al., 2010***; ***Sayal et al., 2016***). Indeed, while Dorsal is the sole maternal nuclear-localized input specifying dorsoventral position in *Drosophila*, it rarely acts alone in endogenous enhancers (***Hong et al., 2008***). For example, the interaction of Dorsal with Twist is a classic example of positive cooperativity in development (***Szymanski and Levine, 1995***). Dorsal can also act as a repressor depending on the presence of nearby Capicua binding sites (***Shin and Hong, 2014***). The minimal synthetic enhancers presented here could be used as scaffolds for more complex minimal enhancers incorporating a second binding site for Twist or Capicua, for example.

In conclusion, we have developed a minimal synthetic enhancer system that has shed light on the fundamental assumptions about transcription in development. By engaging in a dialogue between theory and experiment, we have advanced our understanding of how kinetic processes give rise to important features of transcriptional dynamics in the embryo and made progress toward predictive understanding of how regulatory DNA sequence dictates the functional relation between input transcription factor dynamics and output transcriptional activity in development.

## 4 Methods and materials

### 4.1 Plasmids and reporter design

To design our minimal construct (Fig. 3), we placed the 10 bp consensus Dorsal binding site (***Markstein et al., 2002***) upstream of the *even-skipped* core promoter. This enhancer-promoter construct drives the expression of the MS2v5 sequence containing 24 nonrepetitive MS2 loops (***Tutucci et al., 2018***) followed by the *lacZ* coding sequence and the *tubulin* 3’UTR. (***Garcia et al., 2013***).

In addition to the consensus Dorsal binding site (DBS_6.23), we created six enhancers of varying strength by introducing point mutations to the consensus Dorsal binding motif. Some of these binding sites were taken from known validated Dorsal motifs (***Markstein et al., 2002***), while others were generated based on mutations known to decrease Dorsal binding (***Ip et al., 1992***; ***Jiang et al., 1991***). To guide the design of these binding sites, we used an already existing position weight matrix computed with the MEME algorithm (***Ivan et al., 2008***; ***Bailey et al., 2006***) using motifs generated by DNAse I footprinting assays (***Bergman et al., 2005***) and quantified the information content of each base pair using Patser (***Hertz and Stormo, 1999***).

All plasmid sequences used in this study are shown in Table 1 and can be accessed from a public Benchling folder. Injections were carried out by Rainbow inc. or Bestgene inc.

### 4.2 Flies

Reporter plasmids were injected into BDSC 2y line 27388 containing a landing site in position 38F1. Transgene orientation was confirmed by PCR using primers 18.8 (ggaacgaaggcagttagttgt) and Ori-Seq-F1 (tagttccagtgaaatccaagcattttc) binding outside of the 5’ 38F1 *attP* site and the *even-skipped* promoter, respectively. All reporter lines were confirmed to be in the same orientation. All flies used in this study can be found in Table 2.

To generate the embryos used in the experiments shown in all figures except for Figure 4, we crossed 2x Dorsal or 1x Dorsal virgins to males carrying synthetic enhancers. The genotype of 2x Dorsal flies is *yw;Dl-mVenus (CRISPR), MCP-mCherry; Dorsal-mVenus, MCP-mCherry, His2Av-iRFP*. The genotype of 1x Dorsal flies is *yw;dl[1], MCP-mCherry; Dorsal-mVenus, MCP-mCherry, His2Av-iRFP*. Because there does not seem to be a difference in transcriptional activity between the CRISPR knock-in and the transgene Dorsal-mVenus alleles (Fig. S7), we combined the 1x Dorsal and 2x Dorsal data for some enhancers.

MCP-mCherry and His-iRFP were described before by (***Liu et al., 2021***). The Dorsal-mVenus transgene was developed by ***Reeves et al.*** (***2012***).

To generate the Dorsal-Venus knock-in allele we used the CRISPR/Cas9 protocol described by (***Gratz et al., 2015***). We generated a donor plasmid containing the mVenus sequence followed by a stop codon and a 3xP3-dsRed marker flanked by PiggyBac recombinase sites. This insert was flanked by two ≈1 kbp homology arms matching ≈2 kbp surrounding the Dorsal stop codon (plasmid Dl-mVenus-dsRed in Table 1). The Cas9 expressing BDSC line 51324 was injected with the donor plasmid in combination with a plasmid carrying a sgRNA targeting the sequence GTTGT- GAAAAAGGTATTACG in the C-terminus of Dorsal (plasmid pU6-DlgRNA1 in in Table 1). Survivors were crossed to *yw* and the progeny was screened for dsRed eye fluorescence. Several independent lines were established and tested for rescue. The insertion was confirmed by PCR using primers flanking the homology arms OutLHA (ccattaaaacggaaccaagaggtgag) and OutDlRHA (tctaacaatggctc- gatttttgcca). The dsRed eye marker cassette was flipped out of rescuing lines via crossing with a piggyBac recombinase line. The resulting Dorsal-mVenus locus was then resequenced using the same primers.

The data shown in Figure 4 were obtained from embryos laid by *yw;ParB2-eGFP, eNosx2-MCP-mCherry;+* (wild-type Dorsal mothers) or *yw;ParB2-eGFP, eNosx2-MCP-mCherry, dl[1];+* (Dorsal null mothers).

### 4.3 Microscopy

Fly cages were allowed to lay for 90 to 120 minutes prior to embryo collection. Embryos were then mounted on microscopy slides in Halocarbon 27 oil (Sigma-Aldrich, H8773) in between a coverslip and breathable membrane as described in (***Garcia et al., 2013***; ***Bothma et al., 2014***; ***Garcia and Gregor, 2018***).

Confocal microscopy was performed on a Leica SP8 with HyD detectors and a White Light Laser. We used a 63x oil objective, and scanned bidirectionally with a scan rate of 420 Hz and a magnification of 3.4x zoom. We did not use line or frame accumulation. Time-lapse z-stacks were collected with "’10 s frame rate and 106 nm x-y pixel dimensions and 0.5 *µ*m separation between z-slices (7 *µ*M range, 16 slices). x-y resolution was 512x512 pixels. Pinhole was set to 1.0 Airy units at 600 nm. mVenus was excited by a 510 nm laser line calibrated to 5 *µ*W using the 10x objective and detected in a 520-567 nm spectral window. mCherry was excited by a 585 nm laser line calibrated to 25 *µ*W and detected in a 597-660 nm spectral window. To image His2av-iRFP, the 700 nm laser line was set to 10% and detected in a 700-799 nm spectral window. In all channels, detection was performed using the counting mode of the HyD detectors.

All movies were taken at "’50% along the anterior-posterior axis of the embryo.

### 4.4 ParB experiment 2y crosses and microscopy

We created flies with and without functional Dorsal expressing ParB2-eGFP maternally driven by the *nanos* promoter and MCP-mCherry driven by two copies of a minimal *nanos* enhancer to label our locus DNA and nascent mRNA, respectively. In addition, we added a parS sequence followed by a 400 bp spacer (created with SiteOut, ***Estrada et al.*** (***2016***)) to our DBS_6.23 enhancer. We then crossed male flies containing parS-DBS_6.23-MS2 to *yw; ParB2-eGFP; eNosx2-MCP-mCherry; +* females to create embryos that have our locus of interest labeled with eGFP colocalized with transcriptional loci in the MCP-mCherry channel (Fig. 4A and B).

After mounting embryos using the protocol described above in Section 4.3, we used the sequential scanning mode on the Leica SP8 confocal microscope to eliminate bleedthrough from eGFP into the mCherry channel, and imaged at approximately 20 s per stack, half the rate used in other imaging experiments in this study.

### 4.5 Image and time-series analysis

Image analysis was performed in Matlab using the custom pipeline described in ***Garcia et al.*** (***2013***) and ***Lammers et al.*** (***2020***) (this pipeline can be found in the mRNA Dynamics Github repository). Image segmentation was also aided by the Trainable Weka Segmentation plugin in FIJI (***Witten et al., 2016***; ***Arganda-Carreras et al., 2017***). Further analysis of time-series and other data were likewise performed in Matlab. Movies for publication were made in FIJI (***Schneider et al., 2012***; ***Schindelin et al., 2012***).

### 4.6 Measuring Dorsal-mVenus concentration

Dorsal-mVenus concentration was calculated as in (Fig. S5). As shown in the figure, we measured the average mVenus fluorescence intensity in a circle of 2 *µ*m radius at the center of the nucleus in every z-slice of each nucleus. This results in a z-pro1le of 2uorescence values covering the nucleus itself and the cytoplasm below and above it. The reported concentration corresponds to the value at the middle z-plane of each nucleus. To 1nd this plane, we 1t a parabola to the 2uorescence z-pro1le. We use as the nuclear concentration the fluorescence value at the plane corresponding to the fitted parabola’s vertex (Fig. S5B). We then plotted this value over time and selected a single time point for each trace corresponding to the middle of each nucleus’s observed trajectory (Fig. S5B). To determine the background fluorescence in the mVenus channel we imaged flies with the same genotype as 2x Dorsal except for the Dorsal-Venus fusions. We calculated the average nuclear fluorescence in the mVenus channel across nuclear cycle 12 and subtracted this value from our Dorsal-Venus measurements.

### 4.7 Curve 1tting and parameter inference

Curve fitting and parameter inference were performed in Matlab using the MCMCSTAT Matlab package using the DRAM Markov Chain Monte Carlo algorithm (***Haario et al., 2006***). For simplicity, uniform priors were assumed throughout.

## Acknowledgments

We thank Greg Reeves for providing the Dorsal-mVenus and *dl*^1^ 2y lines. We also thank Francois Payre and Philippe Valenti for sharing a ParB2-eGFP plasmid and a 2xIntB2 (aka *parS*) plasmid. We would like to thank Rob Phillips, Jane Kondev, and members of the Garcia lab for their helpful feedback on the manuscript.

H.G.G was supported by the Burroughs Wellcome Fund Career Award at the Scienti1c Interface, the Sloan Research Foundation, the Human Frontiers Science Program, the Searle Scholars Program, the Shurl and Kay Curci Foundation, the Hellman Foundation, the NIH Director’s New Innovator Award (DP2 OD024541-01), and an NSF CAREER Award (1652236). AR was supported by NSF GRFP (DGE 1752814).

## 5 Biological material

**Table 1.**
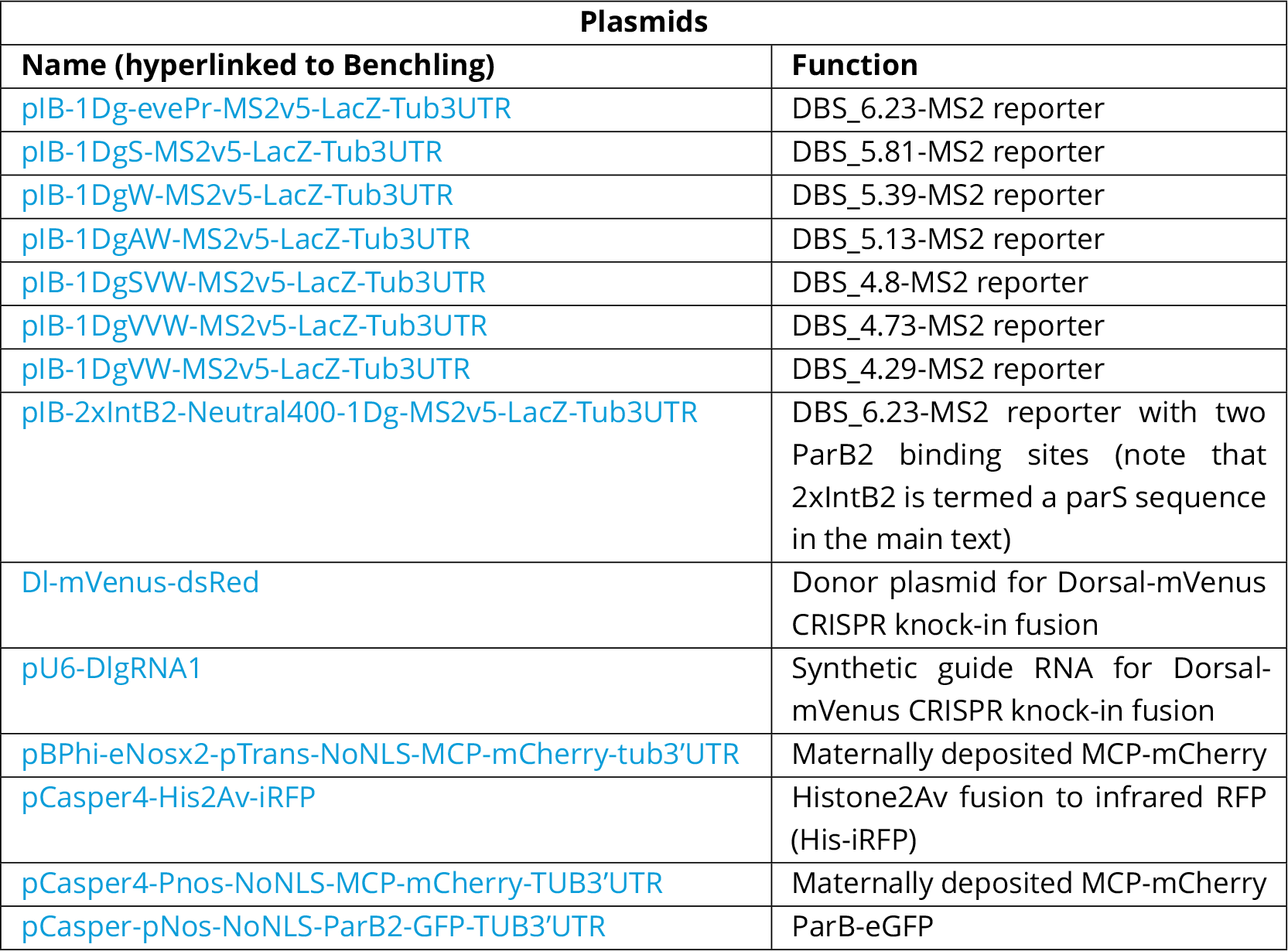
List of plasmids used to create the transgenic 2y lines used in this study.

**Table 2.**
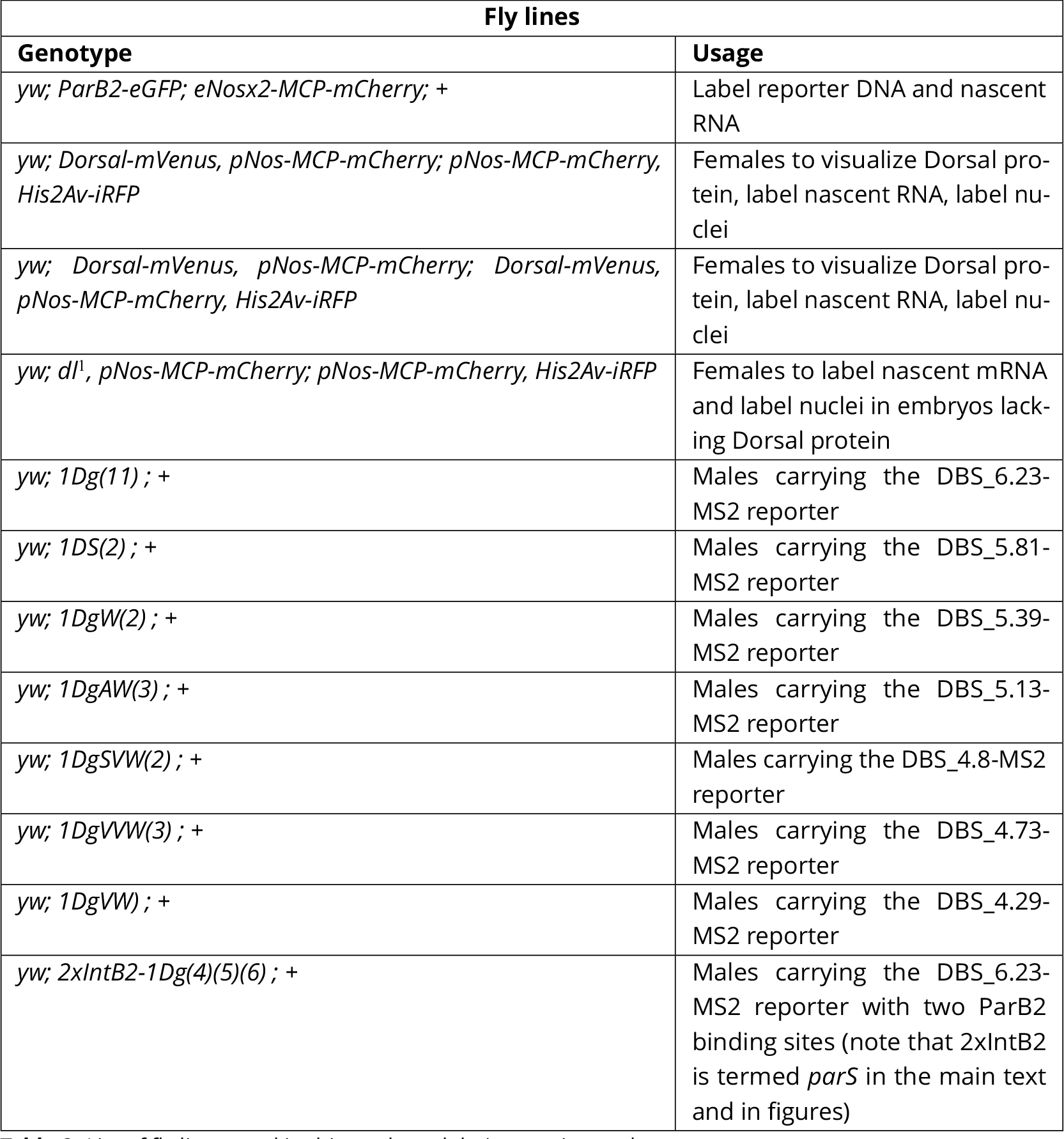
List of 2y lines used in this study and their experimental usage

## Supplementary information

### S1 Appendix

#### S1.1 Calculating the fraction of active loci and the transcriptional onset time by solving the kinetic barrier model

We describe here in detail the method we used to solve kinetic barrier model presented in Section 2.1 and Figure 2A. The problem posed in Figure 5A, namely the time evolution of the probability of nuclei occupying a discrete number of consecutive states, can be described by the following system of linear differential equations (also known as the ‘master equation’)

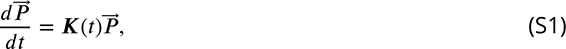

where *P* is a column vector containing the probability as a function of time of each of the states that the system can be in. *K* corresponds to the transition rate matrix containing the rates that dictate the passage from each OFF state to the next and to the final ON state.

For *n* OFF states followed by a ON state connected by irreversible transitions with a rate of *k*(*t*), Equation S1 can be written as

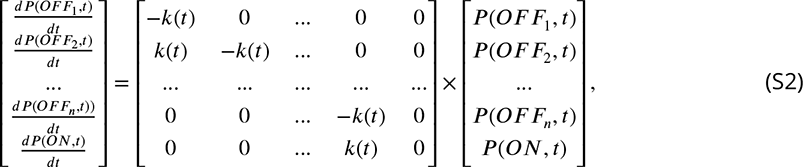

where *P* (*s, t*) indicates the probability of the system being in state *s* at time *t*.

As described in Section 2.4, the transition rate matrix, *K*, is a function of time as a consequence of the assumption that the transition rate between states, *k*, depends on the time-varying Dorsal concentration. In our model, *k* is given by

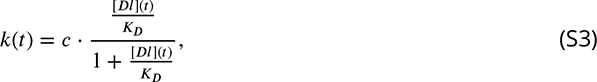

where *K_D_* is the Dorsal binding dissociation constant and *c* is a rate constant. If *k* were a constant, then the system of equations describing transcriptional dynamics could be solved analytically. However, because *k*(*t*) depends on the empirical Dorsal-mVenus fluorescence dynamics, which does not have a concrete functional form, solving the system in Equation S2 becomes analytically intractable. Thus, in order to obtain the probability of each state as a function of time, *P* , and calculate the fraction of active loci and the mean transcription onset times, we solve the system in Equation S2 numerically for a given number of *n* OFF states. Specifically, at each time step *dt*, we calculated how the probability of each state changes with respect to the previous time step.

To calculate *P* (*s, t*) we need to consider the previous time step *t* − 1 and take into account three possible scenarios:

1. Loci that were already in state *s* at time *t* − 1 and stay in this state at time *t*.

2. Loci that were in state *s* − 1 at *t* − 1 that transition into state *s* at time *t*.

3. Loci that were in state *s* at time *t* − 1 that leave this state by transitioning to the next state *s* + 1 at time *t*.

The likelihood of a locus jumping from one state to the next at time *t* during an arbitrarily small time window of *dt* is given by the transition rate *k*(*t*) × *dt*. As a result, the probability of the promoter locus being in state *s* at time *t* can be calculated as

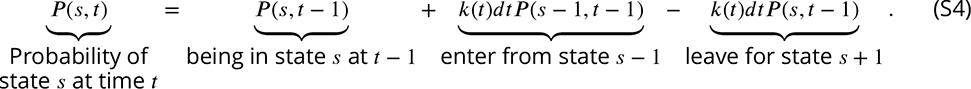

It is clear that, for *s* = 1, *P* (*s* − 1*, t* − 1) = 0, since there is not a previous state from which loci can enter the 1rst OFF state. Similarly, since promoters cannot leave the final ON state once they have entered it, *P* (*n* + 2*, t* − 1) = 0 for *n* OFF states.

To obtain the fraction of active loci, we initialize the system to *P* (*s* = 1*, t* = 0) = 1 and calculate

*P* (*s* = *n* + 1*, t* = *T* /*dt*), where *T* is the duration of the transcriptional window such that

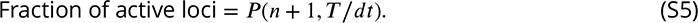

To obtain the mean transcriptional onset time, we calculate the expected value IE[*onset*] of the time to reach the final *n* + 1 state before the end of the transcriptional time window at *t* = *T* . From the de1nition of expected value,

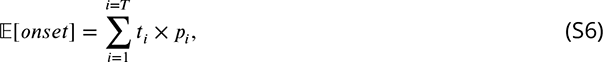

where *t_i_* indicates a given onset time and *p_i_* the probability of loci having that specific onset time. Note that the sum only runs until the end of the transcription time window *T* , as loci that will remain inactive for the duration of the nuclear cycle should not be considered in our calculation of the mean transcriptional onset time. This means that *p_i_* is a normalized probability, calculated only amongst loci that turn on before time *T* such that

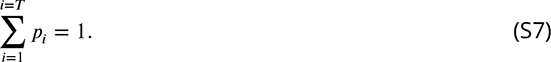

In terms of the system described in Equation S4, the probability *p_i_* of loci reaching the ON state *n*+1 at time *t_i_* is

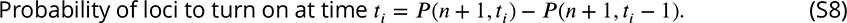

And the normalized probability *p_i_* of loci reaching the ON state *n*+1 at time *t_i_* among loci that reach it before *T* is

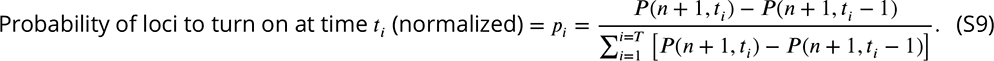

Replacing *p_i_* in Equation S6 with its definition in Equation S9, we arrive at the formula for the mean transcriptional onset time

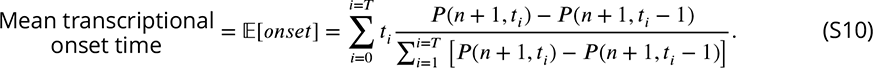

Note that the solutions for the fraction of active loci (Eqn. S5) and their mean transcription onset time (Eqn. S9) ultimately depend on the Dorsal concentration over time [*Dl*](*t*) as they determine *P* (*t, n*). Hence, to generate predictions that can be directly compared to our live-imaging measurements, we need to solve these equations accounting for the Dorsal-mVenus 2uorescence dynamics that determine [*Dl*](*t*).

#### S1.2 Relating MS2 signal to the statistical mechanical model

In order to understand how the maximum MCP-mCherry fluorescence of a locus relates to the average RNAP loading rate, a model for the fluorescence trajectory during a nuclear cycle is required. We start by assuming that RNAP molecules begin loading at a time *t*_0_ into the nuclear cycle and continue to load at a constant rate proportional to *R*, as shown in Equation 2 (*R* = *R_max_* · *p_bound_* ) and step (1) in Figure S1. The observed signal increases linearly until the 1rst polymerase terminates transcription. At this point, the signal plateaus at the value *f_max_* because polymerase molecules continue to be loaded onto the gene at a constant rate while simultaneously terminating at the same rate at the end of the gene (Fig. S1, step (2)). We note that, in this model, initiation halts at step (3), leading to a decrease in 2uorescence as elongating polymerases 1nish transcribing. Note that this step is not accounted for in any analyses or models in this study.

**Figure S1.**
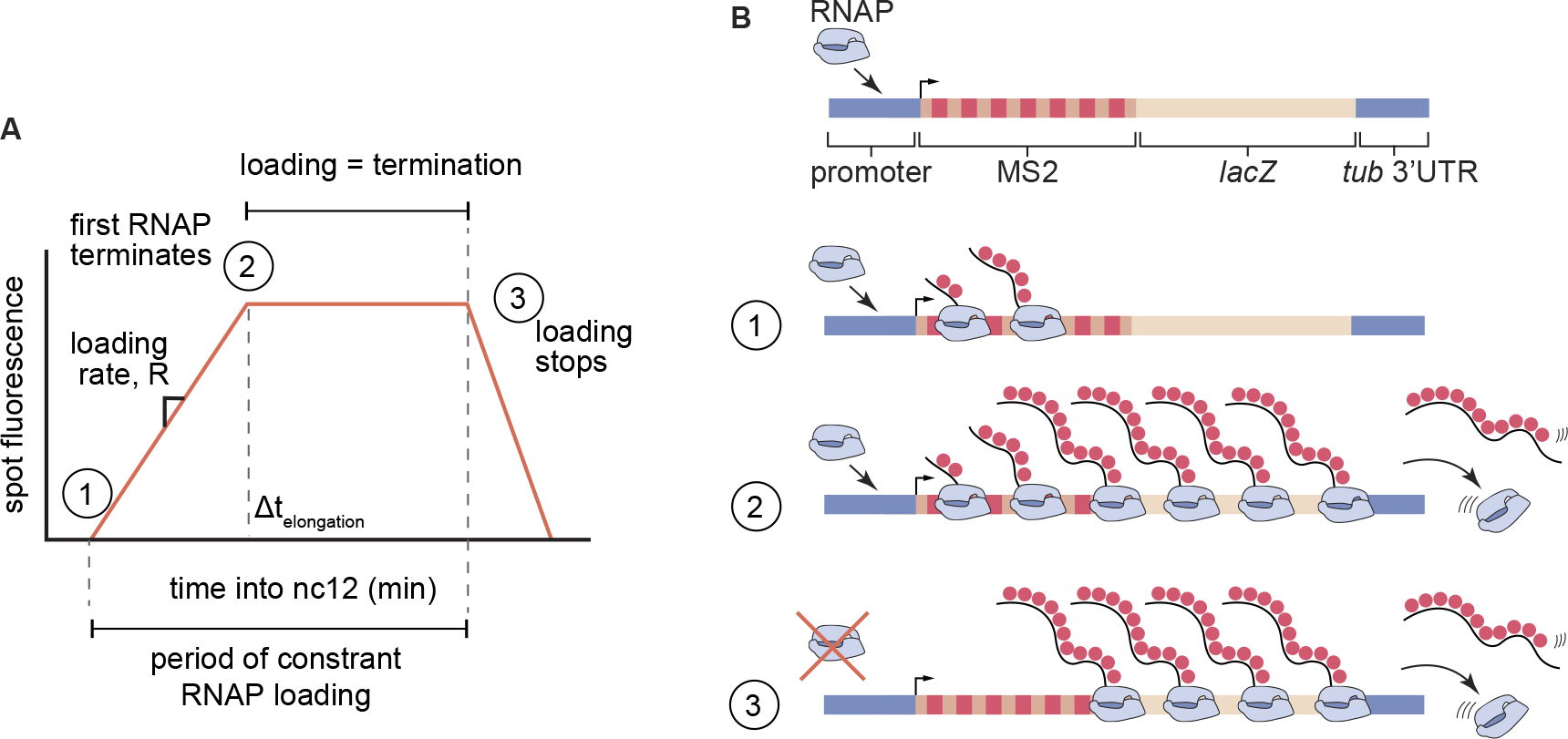
Trapezoid model of transcription dynamics during early embryonic nuclear cycles in *Drosophila*. **(A)**Depiction of a piece-wise linear approximation to average measured fluorescence of loci as a function of time during nuclear cycle 12. In step (1), RNAP molecules are loaded on to the gene at an average constant rate, *R*. After the first RNAP terminates transcription at time fl*t_elongation_*, initiation and termination balance each other out, leading to a constant fluorescence value (step (2)). In step (3), initiation ends, causing the observed fluorescence to monotonically decrease. **(B)** Schematic of the RNAP loading behavior at each step in (A).

Given this model, the maximum fluorescence observed in a trace is given by

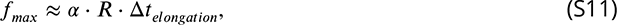

where *R* is the loading defined in Equation 2, and *α* is the instantaneous 2uorescence per mRNA molecule that we estimate in Section S1 .3. As a result, the maximum fluorescence is proportional to the loading rate, namely

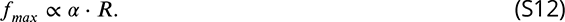

Thus, we now have an expression for *f_max_* that enables us to relate our measurements to the thermodynamic model’s prediction for *R*, the RNAP loading rate (Fig. 2E).

#### S1 .3 MS2 Calibration

To estimate the fluorescence detection threshold in our system, we calibrated the MCP-mCherry signal to single molecule fluorescence *in situ* hybridization (smFISH) data from ***Garcia et al.*** (***2013***). This calibration is based on the fact that, to produce one mRNA molecule, RNAP has to spend a defined amount of time on the reporter thus contributing to the integrated spot 2uorescence. We de1ne *a* as the 2uorescence of one RNAP molecule bearing a labeled nascent RNA and fl*t_elongation_* as the time RNAP spends on the reporter gene to synthesize one mRNA molecule (Fig. S2A). Then, the integrated spot fluorescence corresponding to the production of one mRNA molecule, *p*, is

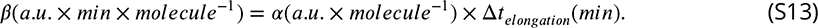

From the de1nition of *β* above, it follows that the integrated fluorescence of a spot over time corresponds to the total number of mRNA molecules produced by that locus in that period (Fig. S2A). Using smFISH, ***Garcia et al.*** (***2013***) measured the mean number of mRNA molecules produced per nucleus by a P2P-MS2 reporter transgene during nuclear cycle 13 as a function of anterior-posterior position (Fig. S2B). To compare these data with the measurements obtained from our imagingsetup, we imaged the same reporter using 2x Dorsal flies and calculated the mean integrated spot 2uorescence across all nuclei as a function of position along the anterior-posterior axis (Fig. S2B). We plotted these two measurements against each other and fitted the data to a line going through the origin (Fig. S2C). The slope of this line indicates *β*, the integrated spot 2uorescence corresponding to a single produced mRNA molecule.

With this 2uorescence calibration factor in hand, we can now estimate *α*, the spot 2uorescence corresponding to a single RNAP molecule attached to one nascent mRNA molecule with 24 MS2 loops. We can estimate fl*t_elongation_* by invoking the elongation rate of RNAP in the 2y embryo, *v_elon_* , and the length of our reporter, *L*, such that

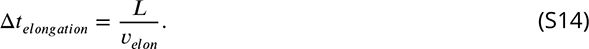

Using this expression for fl*t_elongation_*, we can solve for *α* in Equation S13 to obtain the 2uorescence of a single RNAP molecule given by

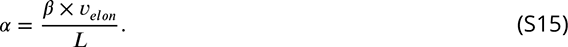

We next replace *L* by the length of our reporter transgene, 5.2 kbp. In addition we replace *v_elon_* by a previously experimentally measured value of 1.5 ± 0.14 kbp/min (***Garcia et al., 2013***), and *β* by the calibration factor shown in Figure S2C. We then arrive at

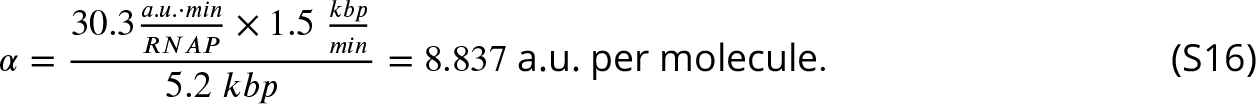

Note that *v_elon_* and *β* have an associated error that leads to uncertainty in the calculation of *a*. Propagating these errors results in an uncertainty of 0.046 a.u. per RNAP, or approximately 14%. This uncertainty should be viewed as an underestimate since, for example, we are not accounting for embryo-to-embryo variability in the accumulated mRNA measured by microscopy or smFISH.

Using this calibration factor, we can now determine the detection threshold of our experimental setup in terms of absolute number of RNAP molecules. One way of determining this threshold is by comparing the mean fluorescence of the dimmest spots with the magnitude of their corresponding background 2uctuations. If these values overlap, then it is not possible to determine with certainty whether a spot correspond to actual signal or to background. This approach reveals a detection threshold of ≈ 80 a.u. or ≈ 9 RNAP molecules (Fig. S2D). A second strategy to determining the detection threshold is looking at the fluorescence of the dimmest detected spots. Their average 2uorescence indicates the value under which no reliable detection is possible. This analysis reveals a detection limit of ≈ 54 a.u. or ≈ 6 RNAP molecules (Fig. S2E). These values for our detection limit using MCP-mCherry are on the order of twice the limit determined for similar experiments that used MCP-eGFP or PCP-eGFP (***Garcia et al., 2013***; ***Alamos et al., 2020***), most likely due to mCherry being a dimmer fluorophore than eGFP (***Lambert, 2019***).

Finally, in the main text (Section 2.5), we estimated the maximum 2uorescence corresponding to the basal level of RNAP molecules on our reporter constructs (Section 2.5). We include here details of the calculation. Since the length of the coding region of our reporter constructs is 5.2 kbp, and the footprint of RNA Polymerase II is 40 bp (***Selby et al., 1997***), 130 RNAPs can 1t on the gene at any given time. Since we estimate the maximum fluorescence corresponding to basal transcription to be ≈ 20 RNAP molecules (Section 2.5), the reporter is 20/130≈ 15% saturated by RNAPs.

**Figure S2.**
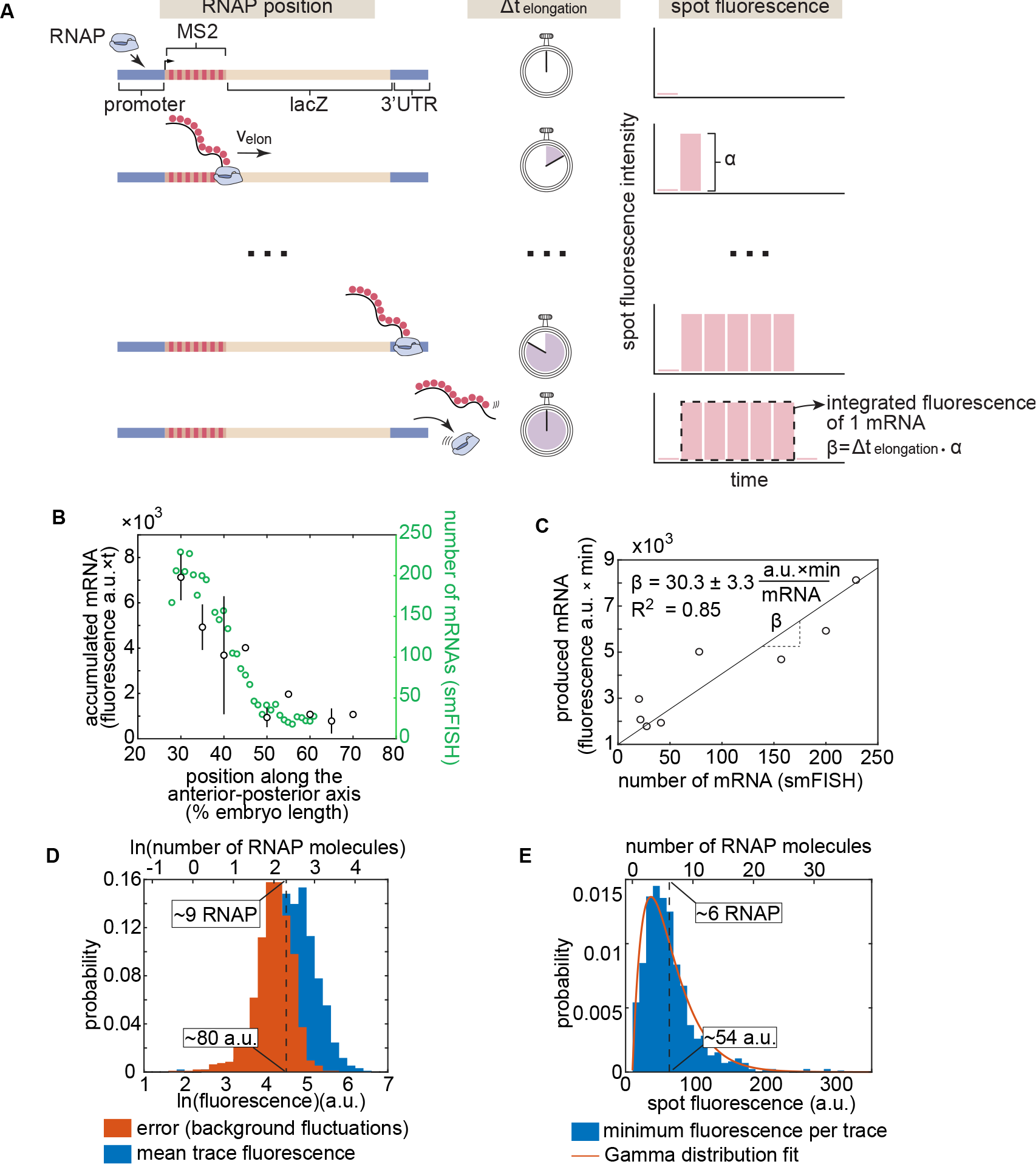
Absolute calibration of MS2 using single molecule FISH. **(A)** Schematic showing that the integrated spot fluorescence corresponding to the production of one mRNA, *β*, is equal to the fluorescence of a single RNAP molecule, *α*, multiplied by the time it spends on the gene, fl*t_elongation_*. **(B)** Mean accumulated mRNA per nucleus (in nuclear cycle 13) based on the integrated MS2 fluorescence of P2P-MS2 employing the imaging conditions used for our reporter data (N = 6 embryos) compared to the number of mRNA molecules per nucleus produced in nuclear cycle 13 as reported by single molecule FISH by ***Garcia et al.*** (***2013***). **(C)** Scatter plot showing data from (B) corresponding to the same anterior-posterior bin. The solid line shows the best linear 1t to all data points. The slope error corresponds to the standard error of the 1t. The error in the fluorescence per RNAP is the propagated standard error taking the errors in elongation rate and calibration slope into account as described in this section’s text. **(D)** Histograms of mean trace fluorescence in all particles across all experiments and the error in the fluorescence of these particles as reported by 2uctuations in the 2uorescence background. Because the spot fluorescence was obtained by integrating over three slices, the corresponding error was propagated by multiplying the error from one slice (using the method described in (***Garcia et al., 2013***)) by 3. The dashed line indicates the center of where the two distributions overlap, suggesting a detection limit of approximately 9 RNAP molecules. **(E)** Histogram of the minimum spot fluorescence per trace across all experiments. The dashed line indicates the mean of the distribution, suggesting a detection limit of approximately 6 RNAP molecules. Note that in (D) and (E) the top x-axis is expressed in terms of absolute number of RNAP molecules using the calibration from (C). A best 1t to a Gamma distribution is shown in red for ease of visualization.

#### S1.4 Measuring transcriptional onset times

We measured the time at which each locus turns on by determining the first time point where a spot was detected. To make this possible, we needed a reliable way to estimate t = 0 which corresponded to the beginning of the nuclear cycle.

Typically, fluorescently labeled histone is used to determine the timing of anaphase (***Garcia et al., 2013***). However, only a small fraction of our embryos had measurable levels of visible Histone-iRFP, most likely due to embryo-to-embryo variability and the low density of DNA in the nucleus in nuclear cycle 12 (compared to later nuclear cycles when His-iRFP is more visible). When the Histone-iRFP signal was insufficient to determine anaphase, we relied on the Dorsal-mVenus channel. As we describe below, just like Histone-iRFP, the nuclear Dorsal fluorescence also shows a characteristic pattern during mitosis.

To precisely determine which features of the Dorsal-mVenus channel to use for mitosis timing, we imaged Dorsal-mVenus and Histone-RFP—which, as opposed to Histone-iRFP, can be consistently detected—simultaneously (Fig.S3). This exercise showed that the edges of nuclei become less well defined as they enter mitosis and then elongate at the beginning of anaphase (Fig. S3). In this way, we could identify precise anaphase frames in movies with no visible Histone-iRFP. Despite using this method, we still estimate that there may be a 2-3 frame error (i.e. 20-30 s) in our determination of anaphase. Thus, this error is *<* 20% of the measured period of transcriptional activity within nuclear cycle 12 ("’ 3 min).

**Figure S3.**
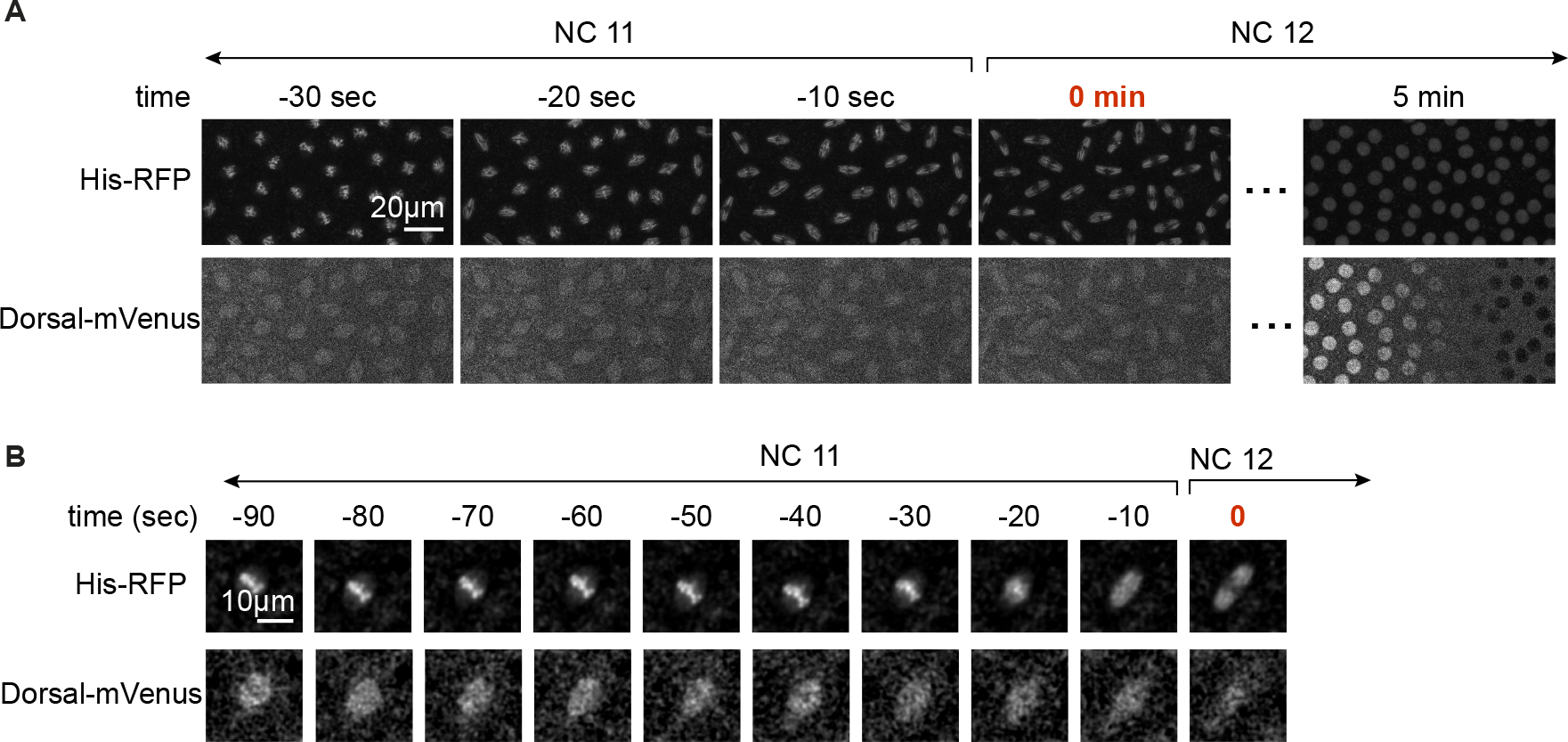
Using the Dorsal-mVenus channel to determine the timing of mitosis. **(A)** Visual comparison of nuclei in the field of view of Histone-RFP and Dorsal-mVenus channels during nuclear division. **(B)** Same as (A), but zoomed into a single nucleus. In (A) and (B), t = 0 min in red text corresponds to anaphase.

#### S1.5 Kinetic barrier 1ts with a different functional form of the transition rate k

In the main text, we hypothesize that the transition rate between OFF states and between the last OFF state and the ON state is proportional to Dorsal occupancy (Eqn. 1). Here, we show that another functional form for *k* in the kinetic barrier model can only partially recapitulate the fraction of active loci and transcriptional onset times for each of our enhancers. This functional form is motivated by the idea that Dorsal could catalyze a change in the promoter (e.g. opening of chromatin) in a manner dependent on the speed of its first occurrence of binding rather than its equilibrium occupancy. Specifically, inspired by (***Eck et al., 2020***), we posit that

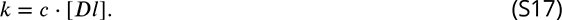

In this alternate model, we assume that the Dorsal binding site affinity dependence is wrapped up into the *c* parameter. Thus, we 1t each enhancer using a distinct value of *c*. As can be seen in Figure S4, this alternate model cannot 1t the data as well as when *k* is assumed to be proportional to the Dorsal occupancy as described in the main text and in Figure 5. Specifically, this functional form is less capable of recapitulating the saturation plateau of the fraction of active loci at high Dorsal concentrations.

**Figure S4.**
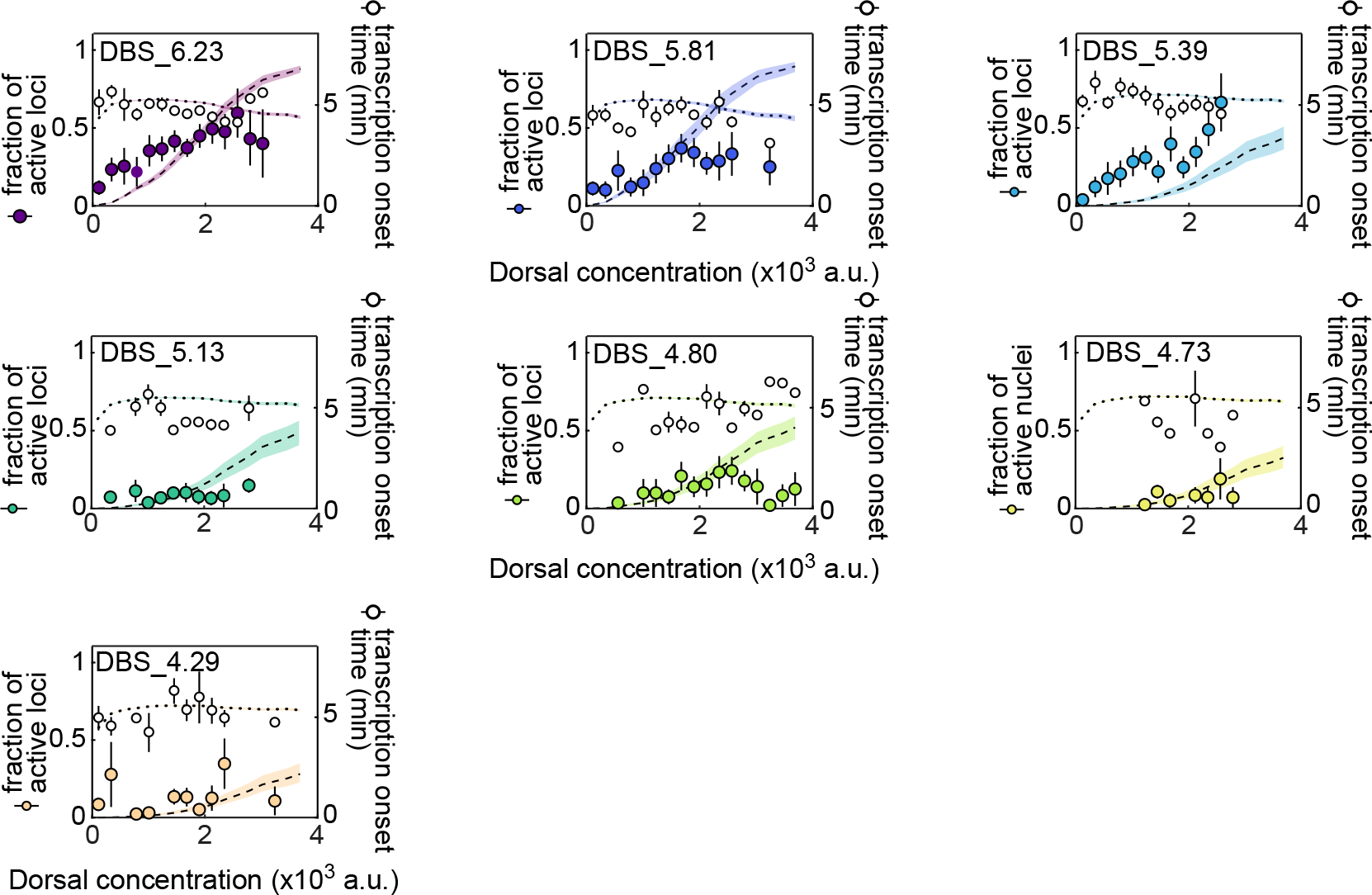
Fits to kinetic barrier model using k = c · [DI]. Data and model 1ts for the fraction of active loci (left y-axis) and mean transcription onset time (right y-axis) for each enhancer. Empty black circles correspond to the experimentally observed mean transcription onset time. Filled colored circles correspond to experimentally observed mean fraction of active loci. Error bars on observations correspond to the standard error of the mean. Fitted curves are represented as black dashed lines (fraction of active loci) and black dotted lines (mean transcription onset times), which correspond to predictions using median parameter values from the joint posterior distribution. Colored shaded areas indicate the 25%-75% credible interval.

### S2 Supplementary Figures

**Figure S5.**
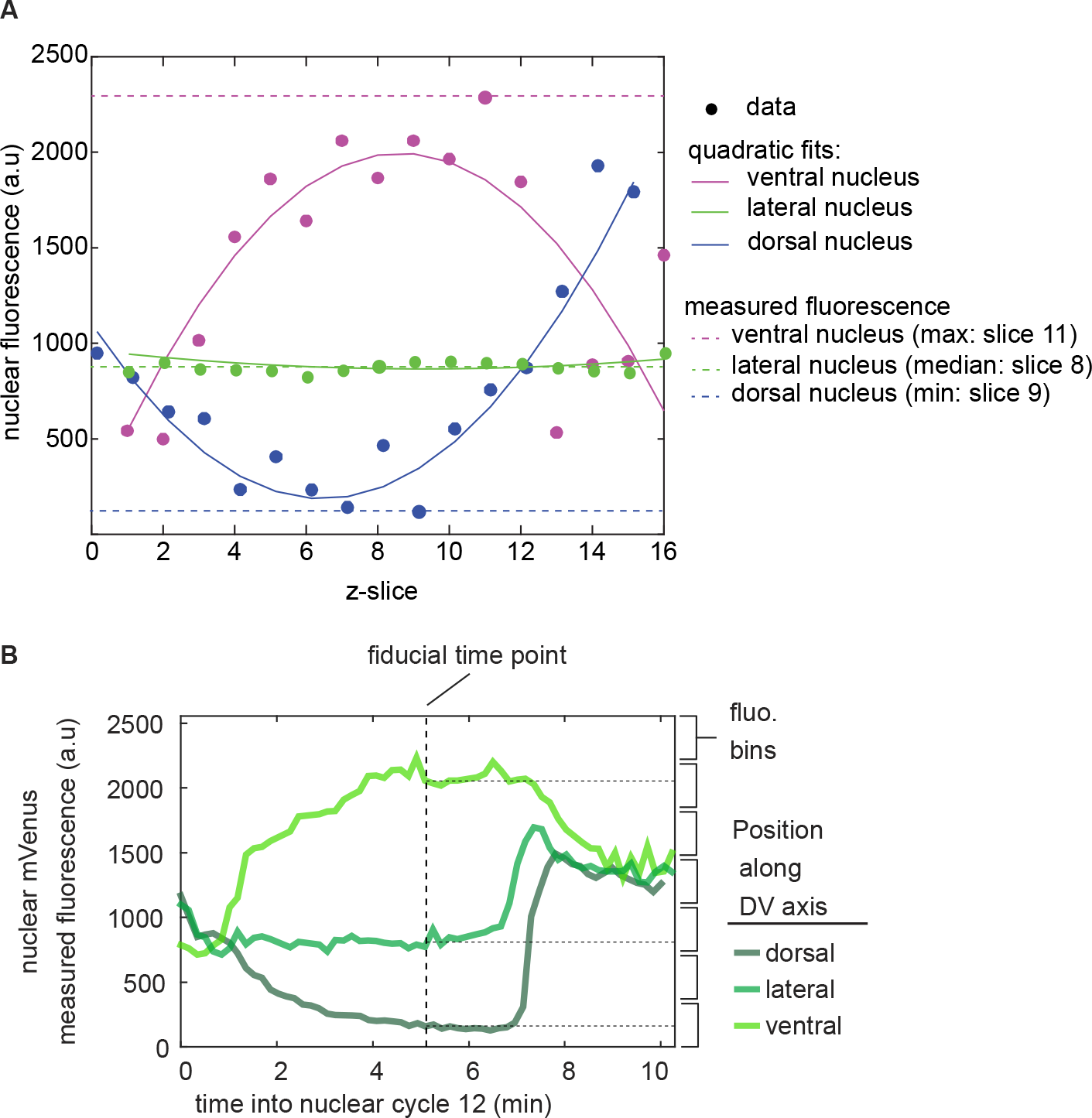
Measuring Dorsal-mVenus nuclear fluorescence across the dorsoventral axis. **(A)** In each frame, the Dorsal-mVenus fluorescence is measured in each z-slice across nuclei. This creates a series of fluorescence values as a function of z-slice (filled circles). z-slices at the top and the bottom correspond to cytoplasmic fluorescence. Thus, in ventral nuclei, the brightest slice is the z-slice corresponding to the best estimate of the true nuclear fluorescence (magenta circles). On the other hand, dorsal nuclei have a lower Dorsal concentration than the cytoplasm, so the darkest slice is a better estimate of the true Dorsal concentration (blue circles). In lateral nuclei, the nuclear fluorescence is similar to that of the cytoplasm (green circles). To identify which z-slice to use for nuclear fluorescence calculations, we 1t the 2uorescence, *f* , overlices, *z*, to a quadratic equation, *f* = *az*^2^ + *bz*, where *a* and *b* are the coefficients of this quadratic equation. Then, we use the value of *a* to determine whether the nucleus is ventral (*a* < -0.5), lateral (-0.5 *a* <0.5), or dorsal (*a* > 0.5). Next, in ventral nuclei, we take the brightest z-slice as the Dorsal-mVenus fluorescence of that frame (dashed horizontal magenta line). In lateral nuclei, we take the median of fluorescence values over z-slices (dashed horizontal green line). In dorsal nuclei, we take the darkest z-slice as the respective frame’s Dorsal-mVenus fluorescence (dashed horizontal blue line). **(B)** Representative time traces of nuclear Dorsal-mVenus fluorescence. To calculate transcriptional activity as a function of Dorsal protein, we sort nuclei into Dorsal concentration bins based on the the Dorsal-mVenus fluorescence at a single fiducial time point halfway through the respective lifetime of each nucleus.

**Figure S6.**
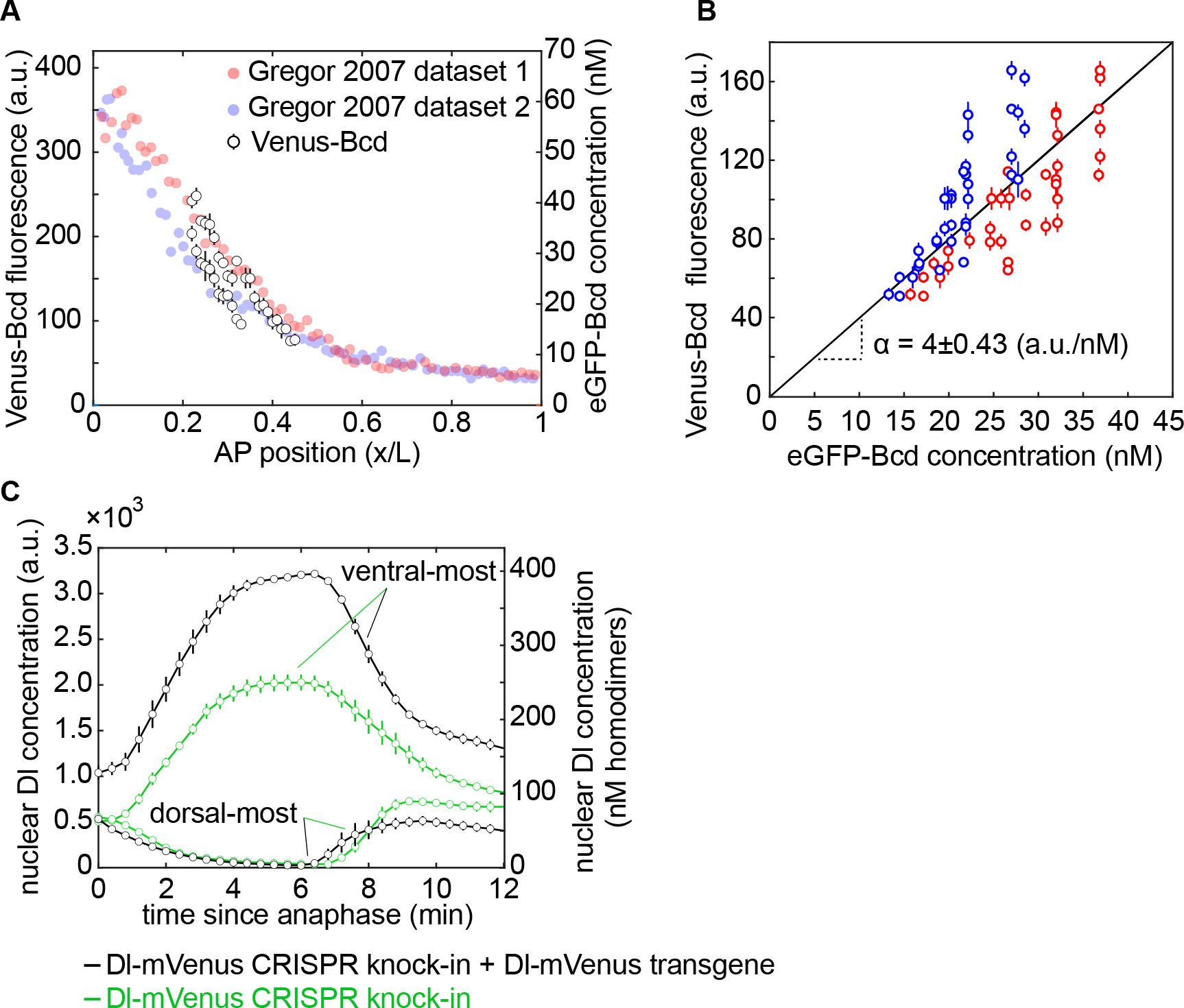
Absolute calibration of Dorsal-mVenus fluorescence using Venus-Bicoid and previously measured eGFP-Bicoid concentration. **(A)** Three embryos derived from *yw;Venus-Bicoid;BcdE1* homozygous mothers were imaged in nuclear cycle 14 using the imaging conditions of our MS2 experiments. The nuclear fluorescence was calculated 15 min into nuclear cycle 14 for cross-comparison with absolute eGFP-Bicoid concentration measurements from Figure 2B of ***Gregor et al.*** (***2007***). We compare the fluorescence values of Venus-Bicoid to the absolute concentration of eGFP-Bicoid along the anterior-posterior axis of the embryo. **(B)** Plot of Venus-Bicoid fluorescence as a function of eGFP-Bicoid fluorescence. Each data point corresponds to the mean ± standard deviation of the fluorescence of all nuclei belonging to the same 1% spatial window along the anterior-posterior axis. These data were compared to two different absolute measurements of eGFP-Bicoid, shown in red and blue. Linear 1t was performed assuming no intercept term since we are estimating a proportionality constant. The slope’s error (*α*) corresponds to the 95% confidence interval. **(C)** Mean and SEM of the Dorsal nuclear concentration in the ventral-most and dorsal-most nuclei across four embryos. 1x and 2x correspond to embryos from homozygous females containing one or two Dorsal-mVenus alleles, respectively. The right y-axis shows the concentration of Dorsal homodimers assuming 6 fluorescence a.u. per mVenus molecule based on (A) and (B).

**Figure S7.**
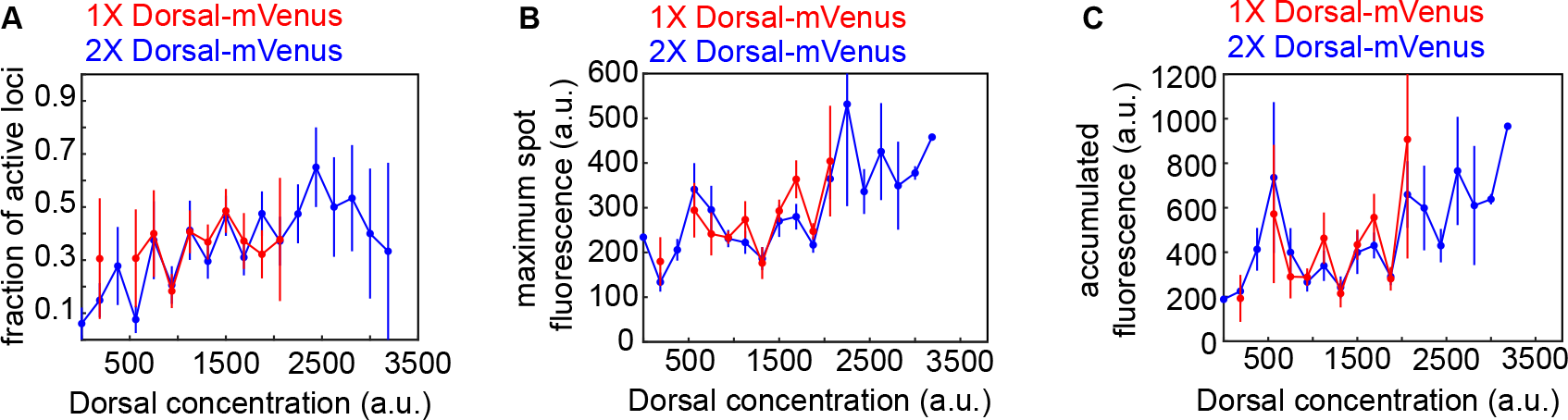
Comparing the activity of the Dorsal-mVenus transgene to that of two copies of Dorsal-mVenus provided by a transgene plus a CRISPR knock-in. For the DBS_6.23 reporter construct, we imaged embryos laid by two different mothers. 1x mothers (red) carry *dl*^1^ (a null Dorsal allele) and a Dorsal-mVenus transgene created by *Reeves et al.* (*2012*). 2x mothers (blue) carry a Dorsal-mVenus CRISPR knock-in and the aforementioned Dorsal-mVenus transgene. Nuclei from these different mothers were binned according to their mVenus fluorescence and different activity metrics were measured for each bin. The two Dorsal-mVenus populations are not different within error such that it is valid to treat embryos laid by these different mothers as equivalent. (Error bars correspond to the standard error across at least three embryos per Dorsal-mVenus fluorescence bin.)

**Figure S8.**
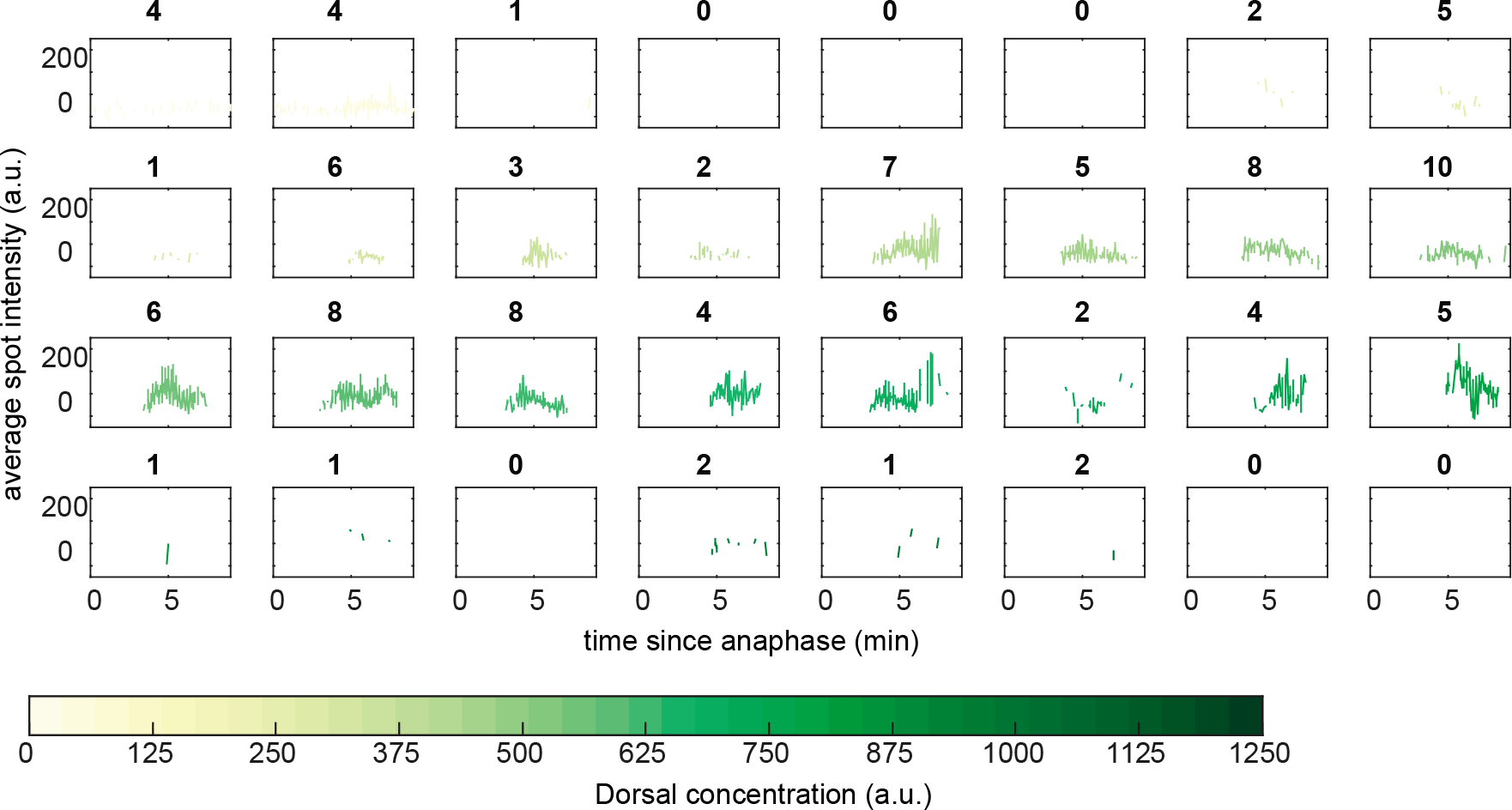
Mean DBS_6.23 transcription spot intensity over time. Mean spot intensity from DBS_6.23 transcription spots over time. Each plot corresponds to a different Dorsal-mVenus concentration as indicated by the legend. The nature of the data makes it challenging to estimate the RNAP loading rate by fitting a line to the initial rise of fluorescence. Bold letters above each plot indicate the number of particles included in each bin.

**Figure S9.**
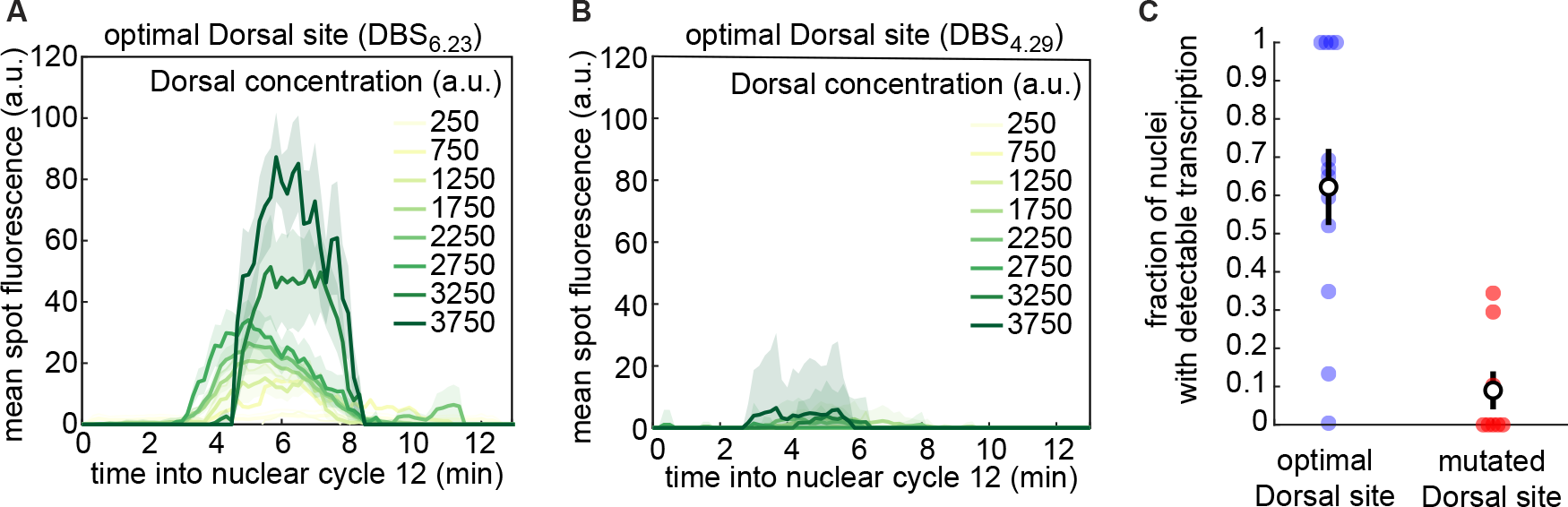
Transcription driven by a minimal Dorsal synthetic enhancer with a mutated Dorsal binding site. **(A,B)** Mean fluorescence over time across all loci in the field of view from an embryo carrying a minimal synthetic enhancer with a **(A)** single optimal and **(B)** a mutated Dorsal binding site. **(C)** Fraction of nuclei in which we detected a transcription spot at any time during the duration of nuclear cycle 12 in nuclei exposed to high Dorsal concentration (2600–3200 a.u.) within the field of view. Filled circles correspond to individual embryos. Black circles show the mean across all embryos. Shaded areas in (A) and (B) and error bars in (C) correspond to the standard error of the mean.

**Figure S10.**
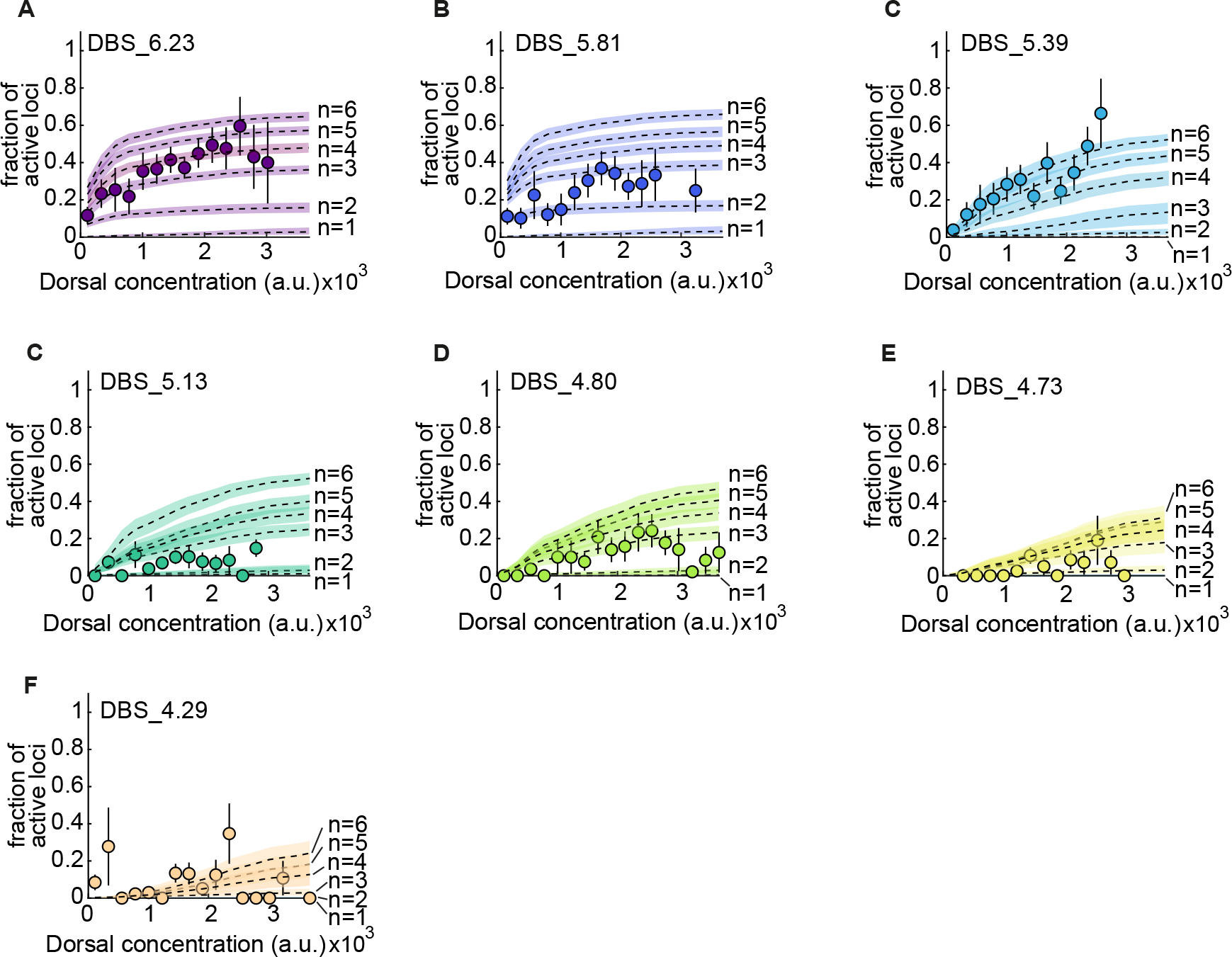
Fits of the kinetic barrier model to the fraction of active nuclei using different numbers of transitions,*n*. **(A)** Mean fraction of active loci as a function of Dorsal concentration in the DBS_6.23 enhancer. Dashed lines show model 1ts using different number of OFF states *n* = 1, 2, 3, 4, 5, and 6, corresponding to predictions using median parameter values from the joint posterior distribution. Fits are performed simultaneously across all enhancers with the value of *c* being shared and the value of *K_D_* being allowed to vary across enhancers. The shaded areas indicate the 25%-75% credible interval. **(B-F)** Same as (A) for the rest of minimal synthetic enhancers. Error bars in (A)-(F) correspond to the SEM taken over N > 3 embryos containing 3 or more nuclei in a given bin.

**Figure S11.**
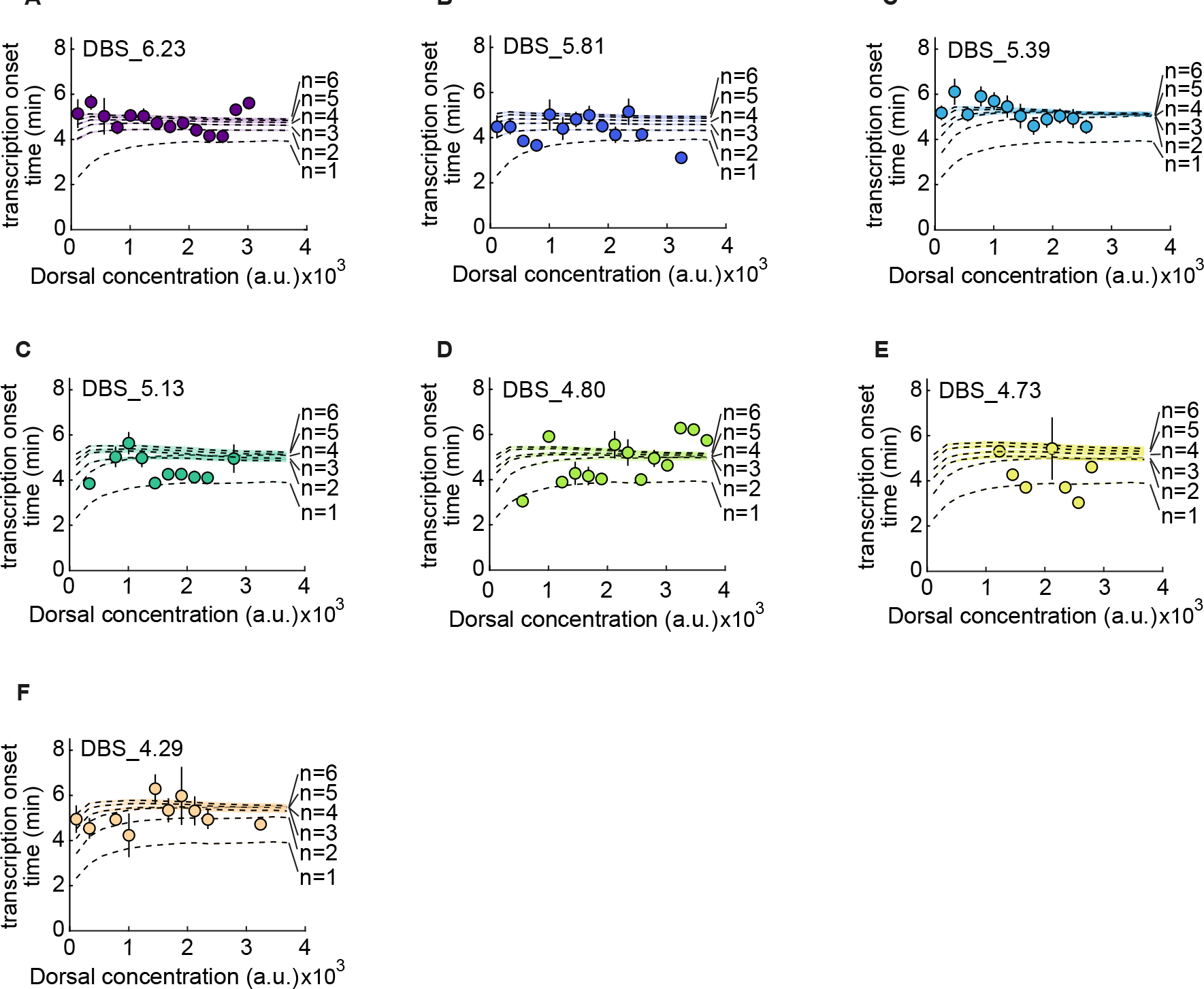
Fits of the kinetic barrier model to the transcription onset times using different numbers of transitions,*n*. **(A)** Mean transcription onset time as a function of Dorsal concentration in the DBS_6.23 enhancer. Dashed lines show model 1ts using different number of OFF states *n* = 1, 2, 3, 4, 5, and 6, corresponding to predictions using median parameter values from the joint posterior distribution. Fits are performed simultaneously across all enhancers with the value of *c* being shared and the value of *K_D_* being allowed to vary across enhancers. The shaded areas indicate the 25%-75% credible interval. **(B-F)** Same as (A) for the rest of minimal synthetic enhancers. Error bars in (A)-(F) correspond to the SEM taken over N > 3 embryos containing 3 or more nuclei in a given bin.

### S3 Supplementary tables

**Table S1.** Inferred parameters from kinetic barrier model 1ts in Figure 5. Each *K_D_* has units of a.u..

| Parameter | Mean | Std. Dev. |
| --- | --- | --- |
| $c$ (min<sup>-1</sup>) | 0.55 | 0.037 |
| $K_D$(DBS_6.23) | 210 | 85 |
| $K_D$(DBS_5.81) | 150 | 52 |
| $K_D$(DBS_5.39) | 980 | 450 |
| $K_D$(DBS_5.13) | 870 | 360 |
| $K_D$(DBS_4.80) | 870 | 340 |
| $K_D$(DBS_4.73) | $1.5 \times 10^3$ | 680 |
| $K_D$(DBS_4.29) | $3.7 \times 10^3$ | $3.1 \times 10^3$ |

**Table S2.** Inferred parameters from 1ts of the thermodynamic model to the RNAP loading rates measured in Figure 6. *R_max_* and *K_D_* each have units of a.u., while the remaining parameters are unitless.

| Parameter | Mean | Std. Dev. |
| --- | --- | --- |
| $R_{max}$ | 510 | 190 |
| $K_D$(DBS_6.23) | $6.3 \times 10^3$ | $5.5 \times 10^3$ |
| $K_D$(DBS_5.81) | $5.4 \times 10^4$ | $2.6 \times 10^4$ |
| $K_D$(DBS_5.39) | $4.3 \times 10^4$ | $2.7 \times 10^4$ |
| $K_D$(DBS_5.13) | $3.9 \times 10^4$ | $2.7 \times 10^4$ |
| $K_D$(DBS_4.80) | $6.7 \times 10^4$ | $2.2 \times 10^4$ |
| $K_D$(DBS_4.73) | $6.6 \times 10^4$ | $2.3 \times 10^4$ |
| $K_D$(DBS_4.29) | $6.8 \times 10^4$ | $2.2 \times 10^4$ |
| $\omega$ | 14 | 23 |
| $P/K_P$ | 0.65 | 0.23 |

### S4 Supplementary videos

For better quality of visualization, we recommend downloading these videos.

- Video S1. **DBS_6.23 confocal movie.** Confocal microscopy movie taken on the ventral side of a developing 2y embryo (*yw; MCP-mCherry, Dl-mVenus(CRISPR) / DBS_6.23-MS2; MCP-mCherry, Dl-mVenus, His-iRFP / +*) during nuclear cycle 12. Left: Dorsal-mVenus; Right: MCP-mCherry.
- Video S2. **ParB experiment confocal movie.** Confocal microscopy movie taken on the ventrolateral side of a developing 2y embryo (*yw; ParB-eGFP, MCP-mCherry / intB2-DBS_6.23- MS2; +*) during nuclear cycle 12. Left: ParB-eGFP; Right: MCP-mCherry.

